# Stromal Gasdermin D-mediated Pyroptosis Drives Maladaptive CD4⁺ T-cell Remodeling in Tet2-Deficient Hematopoiesis

**DOI:** 10.64898/2026.05.01.722102

**Authors:** Kehan Ren, Xu Han, Ermin Li, Pan Wang, Honghao Bi, Wenjie Cai, Inci Aydemir, Ching Man Wai, Hongshen Niu, Jing Yang, Yijie Liu, Brian Vadasz, Madina Sukhanova, Deyu Fang, Weiguo Cui, Peng Ji

**Affiliations:** Department of Pathology, Feinberg School of Medicine, Northwestern University, Chicago, IL; Robert H. Lurie Comprehensive Cancer Center, Northwestern University, Chicago, IL; Department of Biochemistry and Molecular Genetics, Northwestern University, Chicago, IL

## Abstract

An inflammatory bone marrow microenvironment is increasingly recognized as critical in myeloid disease evolution, yet how stromal inflammation interfaces with adaptive immunity remains poorly defined. Here, we show that stromal pyroptosis drives mutation-specific myeloid expansion by coordinating monocytic remodeling and CD4⁺ T-cell activation. Genetic ablation of gasdermin D in the bone marrow stroma suppressed stromal pyroptosis and attenuated Tet2-deficient myeloid expansion. Tet2 deficiency skewed monocyte and macrophage differentiation toward an activated, antigen-presenting state that interacted with pyroptotic stromal cells to promote expansion of a distinct CD4⁺ T-cell population. These cells expressed canonical T follicular helper markers (*Bcl6*, *Cxcr5*, *Il21*, and *Cd40l*) together with interferon-responsive and tissue-interaction programs, consistent with an inflammation-adapted T_FH_-like state. CD40L produced by these cells reinforced the expansion of Tet2-deficient monocytes and macrophages, establishing a feed-forward stromal-immune circuit. Disruption of this axis through stromal gasdermin D deficiency or CD40L blockade attenuated myeloid expansion in vivo. Consistent with these findings, patients with isolated TET2 loss-of-function mutations exhibited CD4⁺ T-cell skewing and CD40L^+^ T-cell-rich tertiary lymphoid structures in the bone marrow. Together, these data identify a pyroptosis-dependent stromal-immune axis that links early myeloid inflammation to maladaptive remodeling of adaptive immunity and reveals a context-dependent therapeutic vulnerability in Tet2-deficient hematopoiesis.

## Introduction

Bone marrow inflammation driven by dysregulated innate immune signaling and inflammasome activation is increasingly recognized as a key determinant of myeloid neoplasms, including clonal cytopenia of undetermined significance (CCUS) and myelodysplastic syndromes (MDS)^1–5^. While extensive work has established a role for inflammatory cytokines and myeloid-derived innate immune pathways in disease pathogenesis^6–9^, how inflammatory signals arising from distinct genetic lesions interface with the bone marrow microenvironment and adaptive immunity remains poorly understood.

Our previous work established that dysregulated innate immune signaling can drive the pathogenesis of myeloid neoplasms through activating inflammasomes. Using mouse models with dual deletion of Diap1 (mDia1) and miR-146a, genes located within the commonly deleted chromosome 5q region (del(5q)) in human MDS, we demonstrated that increased sensitivity to damage- and pathogen-associated molecular patterns in these mice leads to excessive production of inflammatory cytokines, including IL-6 and TNFα, resulting in ineffective hematopoiesis and age-dependent progression to leukemia^7,10,11^. These studies established bone marrow inflammation as a potent driver of MDS evolution, a finding validated in MDS patients^11^. Notably, many of the most frequently mutated genes in myeloid neoplasms, including TET2, have been independently linked to altered inflammatory signaling^12–16^, suggesting that mutation-specific interactions between hematopoietic cells and the inflammatory microenvironment may play a critical role in shaping disease evolution.

## Results

### Hematopoietic cell-intrinsic gasdermin D-mediated pyroptosis drives myeloid neoplasm pathogenesis in a murine MDS model

To identify inflammatory pathways capable of functionally linking innate immune activation to myeloid disease evolution, we focused on pyroptosis, a lytic form of programmed cell death mediated by inflammasome activation and executed by gasdermin D^17–20^. Pyroptosis has been implicated in inflammatory signaling in myeloid cells^21–23^, yet its causal contribution to the development and maintenance of myeloid neoplasms remains unclear. We first investigated the expression levels of gasdermin D in MDS through a published gene expression dataset of CD34^+^ hematopoietic stem and progenitor cells (HSPCs) from MDS patients^24^, and found it significantly increased across different subtypes of MDS (**Figure 1A**), which was confirmed by immunohistochemical staining of bone marrow from MDS patients (**Figure 1B**). Notably, gasdermin D stains showed a diffuse increase in MDS, suggesting its upregulation in the bone marrow stromal cells as well.

**Figure 1:**
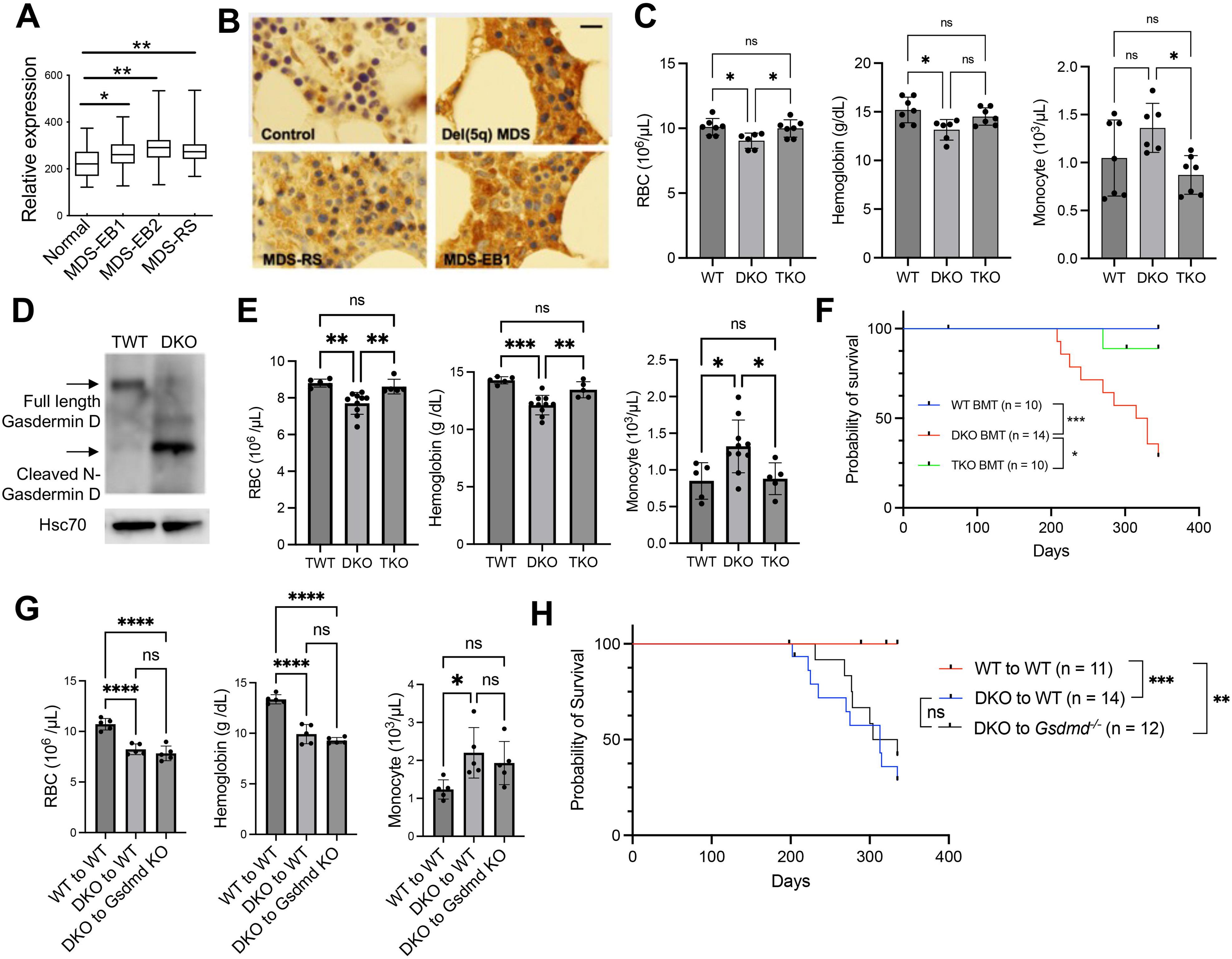
Hematopoietic cell-intrinsic gasdermin D-mediated pyroptosis drives myeloid neoplasm pathogenesis in a murine MDS model. (A) Gasdermin D (GSDMD) mRNA expression in CD34⁺ HSPCs from MDS patients across disease subtypes compared with healthy controls. MDS-EB1: MDS with excess blasts 1; MDS-EB2: MDS with excess blasts 2; MDS-RS: MDS with ring sideroblasts. (B) Representative immunohistochemical staining of gasdermin D in bone marrow biopsies from MDS patients and healthy controls. Scale bar: 100 μm. (C) Red blood cell, platelet, and monocyte count of the indicated mice at 9 months old. Triple wild type (TWT): n = 7; DKO: n = 6; *Gsdmd* TKO: n = 7. (D) Spleen cells from one year old TWT and DKO mice were isolated and applied for a Western blotting assay detecting full length and activated/cleaved N-terminal gasdermin D. (E) Red blood cell count, hemoglobin, and monocyte count of the recipient mice at 9 months post transplantation of the indicated donor bone marrow HSPCs. TWT: n = 5; DKO: n = 10; *Gsdmd* TKO: n = 5. (F) Kaplan-Meier survival analysis of recipient mice transplanted with bone marrow HSPCs from 5-month-old donor mice. (G) Red blood cell count, hemoglobin, and monocyte count of lethally irradiated wild-type (WT) or *Gsdmd^⁻/⁻^* recipients transplanted with bone marrow HSPCs from WT or DKO donors at 9 months post-transplant. n = 5 in each group. (H) Kaplan-Meier survival curves of lethally irradiated WT or *Gsdmd^⁻/⁻^* recipients transplanted with bone marrow HSPCs from WT or DKO donors. All the error bars represent the SD of the mean. Comparisons among multiple groups were evaluated using a 1-way ANOVA. *p<0.05, **p<0.01, ***p<0.001, and ****p<0.0001. ns: not significant.

To directly test the role of gasdermin D in the development and progression of myeloid neoplasms, we crossed constitutive gasdermin D-deficient (*Gsdmd^⁻/⁻^*) mice with mDia1/miR-146a double-knockout (DKO) mice to generate gasdermin D-deficient DKO (TKO) animals. Loss of gasdermin D markedly ameliorated MDS phenotypes in DKO mice, including anemia, thrombocytopenia, and monocytosis (**Figure 1C**). Consistent with functional activation of pyroptosis in this model, gasdermin D was detected in its cleaved form in bone marrow cells from DKO mice (**Figure 1D**). Transplantation of bone marrow lineage-negative (Lin^-^) hematopoietic stem and progenitors (HSPCs) from TKO mice into lethally irradiated wild-type recipients similarly rescued disease phenotypes (**Figure 1E, Supplemental Figures 1A-C**) and significantly extended recipient survival (**Figure 1F**). Given that pyroptosis is known to release inflammatory mediators that can propagate tissue-wide inflammation, we next investigated whether pyroptotic hematopoietic cells could remodel the bone marrow stroma. Indeed, stromal cells from DKO recipients showed marked upregulation of the alarmins S100a8 and S100a9 compared with wild-type controls. Loss of gasdermin D in donor marrow significantly reduced S100a9 expression in both hematopoietic and stromal compartments (**Supplemental Figures 1D-G**), indicating that hematopoietic pyroptosis drives inflammatory stromal remodeling toward an alarmin-high, damage-associated state.

Given that pyroptotic hematopoietic cells induce stromal inflammatory remodeling, we next asked whether stromal pyroptosis could reciprocally contribute to DKO disease pathogenesis. We transplanted DKO HSPCs into either wild-type or gasdermin D-deficient recipient mice that had been lethally irradiated. Stromal gasdermin D deficiency, however, failed to rescue DKO phenotypes or improve survival (**Figures 1G-H**), in contrast to the protective effect of hematopoietic gasdermin D deletion. These results establish that gasdermin D-mediated pyroptosis in hematopoietic cells is the primary cell-autonomous driver of MDS pathogenesis in the DKO model, while stromal pyroptosis is secondarily induced by the malignant hematopoietic compartment.

### Stromal pyroptosis drives Tet2-deficient myeloid expansion

The cell-intrinsic requirement for gasdermin D in the DKO model, where intrinsic inflammasome hyperactivation drives disease, raised the question of whether stromal pyroptosis might instead play a primary, niche-driven role in myeloid disorders characterized by inflammation-responsive mutations. TET2 deficiency provided an ideal system to test this hypothesis. Although TET2-mutant cells exhibit enhanced inflammatory signaling and preferential expansion in inflammatory environments, myeloid disorders occur in only a fraction of humans with TET2 mutation^15,25–31^, suggesting that extrinsic, non-cell-autonomous factors are required for disease onset. However, the cellular source and molecular nature of these niche-derived inflammatory signals within the bone marrow microenvironment remain undefined. We therefore investigated whether stromal pyroptosis could serve as the primary driver of TET2-deficient myeloid expansion. To this end, we transplanted HSPCs from 3-month-old *Tet2^-/-^*(*Vav-iCre*^+^;*Tet2^fl/fl^*) mice into lethally irradiated 3-month-old wild-type or *Gsdmd^-/-^* recipient mice. We monitored complete blood counts and observed that *Gsdmd^-/-^* recipients showed significant attenuation of monocytosis at 3 months post-transplant, preceding the development of anemia and thrombocytopenia that manifested at later stages (**Supplemental Figure 2A**).

By 9 months post-transplant, when *Tet2^-/-^* to wild-type recipients had developed full disease manifestations, *Gsdmd^-/-^*recipients showed complete rescue of both monocytosis and anemia (**Figure 2A**). Flow cytometric analysis of peripheral blood confirmed the normalization of Ly6G^-^Ly6C^+^ monocyte frequencies (**Figure 2B**). The splenomegaly characteristic of *Tet2^-/-^* to wild-type recipients was absent in *Gsdmd^-/-^* recipients (**Figure 2C**). The aberrant splenic cellular compositions, including myeloid (CD11b^+^) expansion and extramedullary erythropoiesis (CD71^+^) that are typical in advanced disease, were also significantly diminished in *Gsdmd^-/-^* recipients (**Figure 2D**). Histopathological examination revealed that *Tet2^-/-^* to wild-type recipients displayed bone marrow with monocytic hyperplasia, effacement of splenic white pulp by expanding myeloid elements, and prominent hepatic myeloid infiltration; all these abnormalities were absent in *Gsdmd^-/-^*recipients, which were histologically indistinguishable from wild-type to wild-type transplant controls (**Figure 2E**).

**Figure 2:**
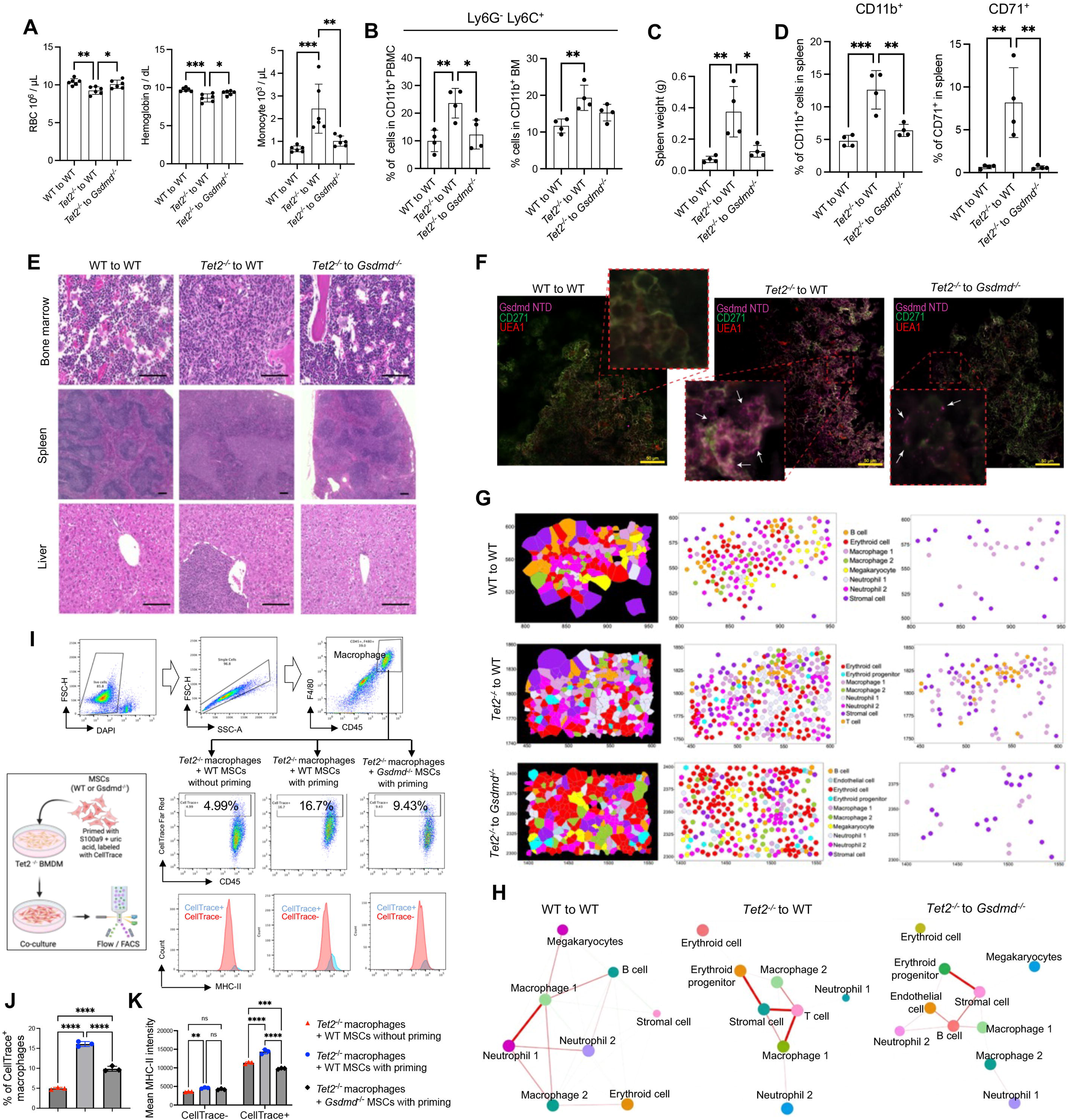
Stromal pyroptosis drives Tet2-deficient myeloid expansion. (A) Red blood cell count, hemoglobin, and monocyte count of lethally irradiated WT or *Gsdmd^⁻/⁻^* recipients transplanted with HSPCs from WT or *Tet2^-/-^* donors at 9 months post-transplant. n = 6 in each group. (B) Flow cytometry analyses of Ly6G⁻Ly6C⁺ monocyte frequencies in CD11b^+^ peripheral blood mononuclear cells (PBMCs) and bone marrow (BM) from indicated transplant groups at 9 months post-transplant. n = 4 in each group. (C) Quantification of spleen weights from the indicated transplant groups at 9 months post-transplant. n = 4 in each group. (D) Flow cytometric analysis of splenic cellular composition showing frequencies of CD11b⁺ myeloid cells and CD71⁺ erythroid precursors from indicated transplant groups. n = 4 in each group. (E) Representative hematoxylin and eosin (H&E) staining of bone marrow, spleen, and liver sections from the indicated transplant groups. Scale bars: 300 μm. (F) Whole-mount confocal imaging of murine femurs showing gasdermin D N-terminal domain (Gsdmd-NTD, magenta) and CD271⁺ mesenchymal stromal cells (MSCs, green) from indicated transplant groups. Arrows indicate punctate membrane-associated Gsdmd-NTD signals indicative of active pyroptosis. Scale bars: 50 μm. (G) High plex Xenium (Xenium 5K) spatial transcriptomic analysis of bone marrow sections from indicated transplant groups. The distributions of monocytes/macrophages relative to stromal cell and T cell populations are highlighted on the right. (H) Cell-cell interaction network analysis demonstrating stromal-macrophage-T cell signaling interactions in each transplant group. The thickness of the connecting lines indicates the strength of the interaction. (I) Schematic of an in vitro co-culture system. The right panels illustrate the increased proportion of CellTrace⁺ *Tet2^⁻/⁻^* macrophages with high MHC-II expression. (J) Quantification of CellTrace⁺ *Tet2^⁻/⁻^* macrophages following co-culture with unprimed or primed wild-type or *Gsdmd^⁻/⁻^* MSCs. n = 3 in each group. (K) Flow cytometric analysis of MHC-II expression on *Tet2^⁻/⁻^* macrophages following co-culture with unprimed or primed MSCs. n = 3 in each group. All the error bars represent the SEM of the mean. Comparisons among multiple groups were evaluated using a 1-way ANOVA. *p<0.05, **p<0.01, ***p<0.001, and ****p<0.0001. ns: not significant.

These findings establish the pyroptotic bone marrow microenvironment as a critical determinant of Tet2-deficient myeloid expansion. To directly visualize pyroptosis within the niche, we applied a whole-mount confocal imaging approach that enables spatial analysis of protein expression within murine femurs. Using an antibody against the pore-forming N-terminal domain of gasdermin D (Gsdmd-NTD), active pyroptosis was identified as punctate membrane-associated signals (**Figure 2F**, arrows). In *Tet2^-/-^* to wild-type recipients, gsdmd-NTD signals were markedly increased not only in donor-derived hematopoietic cells but also in recipient CD271^+^ mesenchymal stromal cells, indicating that Tet2-deficient hematopoietic cells actively remodel the niche by inducing stromal pyroptosis. Strikingly, transplantation into *Gsdmd^-/-^* recipients not only eliminated stromal Gsdmd as expected but also dramatically reduced gasdermin D activation in CD271-negative donor hematopoietic cells (**Figure 2F**). This bidirectional regulation reveals a pyroptotic crosstalk between Tet2-deficient hematopoietic cells and the bone marrow stroma. Mutant hematopoietic cells induce stromal pyroptosis, which in turn amplifies pyroptosis in the hematopoietic compartment, establishing a feed-forward inflammatory loop.

Pyroptotic cell death releases not only inflammatory cytokines but also alarmins and damage-associated molecular patterns (DAMPs), which recruit and activate immune cells at sites of tissue damage^32^. We hypothesize that pyroptotic stromal cells create a niche that preferentially recruits and expands monocytes/macrophages that are characteristic of Tet2-deficient hematopoiesis^15,16^. To test this comprehensively, we performed spatial transcriptomic analysis of cryosectioned bone marrow to map the cellular consequences of stromal pyroptosis. This analysis identified two transcriptionally distinct macrophage populations: macrophage 1, characterized by expression of the monocyte recruitment receptor Ccr2 along with F13a1, Ccl9, and Ms4a6c, consistent with an inflammatory monocyte-derived identity; and macrophage 2, marked by the iron exporter Slc40a1, complement component C1q, and the pattern recognition receptor Tlr4, consistent with a tissue-resident, homeostatic phenotype (**Supplemental Figures 2B-J**). Strikingly, in *Tet2^⁻/⁻^* to wild-type recipients, the inflammatory macrophage population (macrophage 1) was preferentially expanded and spatially clustered in close proximity to pyroptotic stromal cells (**Figure 2G**), establishing a reorganized inflammatory niche architecture that positions these recruited populations for direct cell-cell communication. Interestingly, T cells were also increased and clustered around these macrophage-stromal cell aggregates (**Figure 2G, Supplemental Figures 2B-J**). Cell-cell interaction network analysis revealed a marked enrichment of stromal-macrophage-T cell signaling interactions in *Tet2^-/-^* to wild-type recipients that was absent in both wild-type controls and *Gsdmd^-/-^*recipients (**Figure 2H**). This spatial juxtaposition suggests that pyroptotic stromal cells do not merely release soluble inflammatory signals but also establish direct contact-dependent communication with myeloid cells, promoting their expansion.

To test whether this spatial relationship functionally activates Tet2-deficient myeloid cells, we developed an in vitro co-culture system that recapitulates the stromal-macrophage interaction. CellTrace-labeled mesenchymal stromal cells (MSCs) were primed with S100A9 and uric acid, inflammasome activators known to induce pyroptosis^33,34^, and then co-cultured with *Tet2^-/-^* bone marrow-derived macrophages (**Figure 2I**). We found that the primed wild-type MSCs significantly enhanced *Tet2^-/-^*macrophage uptake of CellTrace-labeled material compared to unprimed controls. This was significantly abolished when *Gsdmd^-/-^* MSCs were used (**Figures 2I, J**), demonstrating a requirement for stromal gasdermin D for the phagocytosis of *Tet2^-/-^* macrophages. Importantly, *Tet2^-/-^*macrophages co-cultured with primed MSCs showed marked upregulation of MHC-II (**Figures 2I, K**), indicating that phagocytosis of pyroptotic stromal material triggers functional reprogramming of *Tet2^-/-^* macrophages toward an antigen-presenting phenotype.

### Stromal pyroptosis licenses the emergence of T_FH_-like CD4^+^ T cells

These data establish pyroptosis in bone marrow mesenchymal stromal cells as a critical driver of Tet2-deficient myeloid expansion. The emergence of MHC-II-high, Tet2-deficient macrophages in the pyroptotic niche suggested a potential bridge to adaptive immunity through engagement of CD4^+^ T cells, a poorly understood process in clonal hematopoiesis. To reveal this, we performed single-cell RNA sequencing analyses of total bone marrow cells after erythroid lysis from wild-type and *Gsdmd^-/-^*recipient mice 9 months post-transplant (**Supplemental Figure 3**). As expected, there was a significant increase in macrophage populations at different levels of maturation in *Tet2^-/-^*to wild-type recipients, which was abolished in *Gsdmd^-/-^* recipients (**Figure 3A**, black circle). Strikingly, consistent with the T-cell expansion observed in our spatial transcriptomic analysis (**Figure 2G**), there was also a marked increase in the T-cell population in *Tet2^-/-^* to wild-type recipients, which was similarly dependent on recipient gasdermin D (**Figure 3A**, red circle).

**Figure 3:**
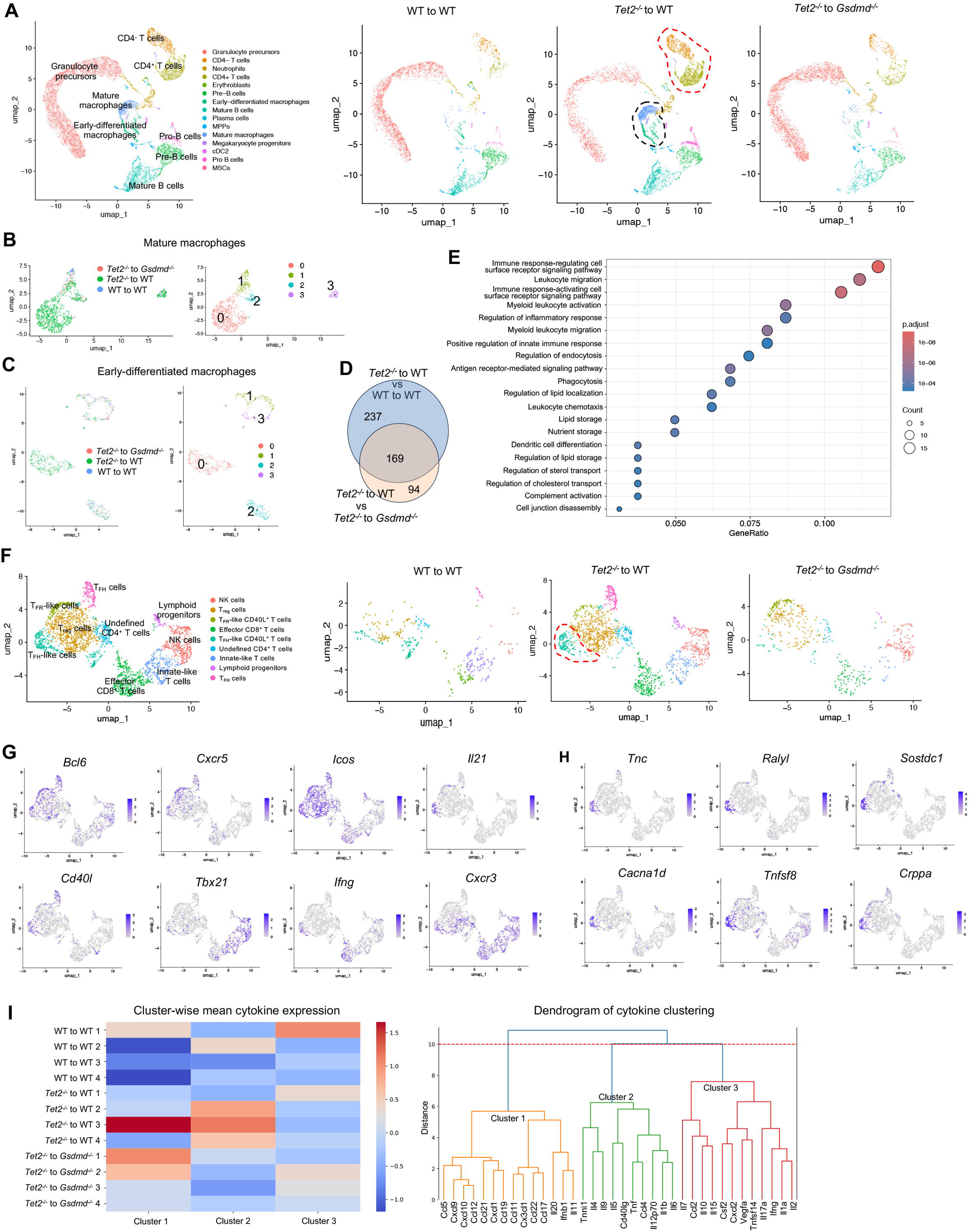
Stromal pyroptosis licenses the emergence of T_FH_-like CD4⁺ T cells. (A) UMAP visualization of single-cell RNA sequencing data from total bone marrow cells (after erythroid lysis) from WT to WT, *Tet2^⁻/⁻^* to WT, and *Tet2^⁻/⁻^* to *Gsdmd^⁻/⁻^* recipients at 9 months post-transplant. The black circle indicates macrophage populations; the red circle indicates T-cell populations. (B-C) UMAP visualization of mature (B) and early differentiated (C) macrophage subpopulations across indicated transplant groups in A. (D) Venn diagram showing overlap of differentially expressed genes in macrophages from *Tet2^⁻/⁻^* to WT recipients compared with WT controls, and genes reversed in *Tet2^⁻/⁻^* to *Gsdmd^⁻/⁻^* recipients. (E) Gene Ontology pathway enrichment analysis of differentially expressed genes in macrophages that are upregulated in *Tet2^⁻/⁻^* to WT recipients and reversed in *Gsdmd^⁻/⁻^* recipients. Dot size represents gene count; color represents adjusted p-value. (F) UMAP visualization of T-cell subpopulations across indicated transplant groups in a, with subclusters annotated by cell type and differentiation state. The T_FH_-like population is highlighted in a red circle. (G-H) Expression of canonical T_FH_ markers (*Bcl6*, *Cxcr5*, *Icos*, *Il21*, *Cd40l*), interferon-responsive genes (*Ifng*, *Tbx21*, *Cxcr3*) (G), and T_FH_-like-specific genes (H) across T-cell subpopulations. (I) Left: Heatmap of cluster-wise mean cytokine expression in bone marrow from WT to WT, *Tet2^⁻/⁻^* to WT, and *Tet2^⁻/⁻^* to *Gsdmd^⁻/⁻^* recipients (n = 4 per group). The color scale represents z-scored expression values. Right: Hierarchical clustering dendrogram of cytokines based on expression patterns across all samples.

Subcluster analyses further revealed that in *Tet2^-/-^* to wild-type recipients, both early-differentiated and mature macrophage populations were markedly expanded relative to controls. This expansion was largely abolished in *Gsdmd^-/-^* recipients, demonstrating that stromal gasdermin D-dependent pyroptosis is required for the sustained accumulation of Tet2-deficient macrophages (**Figures 3B, C**). Consistent with this, differential expression analysis revealed substantial transcriptional remodeling of macrophages in *Tet2^-/-^*to wild-type recipients compared with controls, with a large fraction of these changes overlapping with those reversed in *Gsdmd^-/-^* recipients (**Figure 3D**). Pathway enrichment analysis of these genes revealed that the most significantly enriched pathways were immune response-regulating and immune response-activating cell surface receptor signaling (**Figure 3E**), consistent with the upregulation of MHC-II observed in our co-culture experiments (**Figure 2K**) and suggesting enhanced capacity for antigen presentation to CD4⁺ T cells. Additional enriched pathways included leukocyte migration and chemotaxis, myeloid leukocyte activation, antigen receptor-mediated signaling, and phagocytosis (**Figure 3E**).

These data indicate that stromal pyroptosis not only drives quantitative expansion of Tet2-deficient macrophages but also enforces a qualitatively activated, antigen-presenting macrophage state poised to engage CD4⁺ T cells, providing a mechanistic foundation for downstream adaptive immune remodeling. We therefore next examined how CD4⁺ T cells respond to this pyroptosis-driven myeloid activation by subclustering the T cell population (**Supplemental Figure 4** and **Figure 3F**). In wild-type chimeras, CD4⁺ and CD8⁺ T-cell subsets are distributed across canonical naive, effector, and innate-like states. In contrast, *Tet2^-/-^* to wild-type recipients exhibit a marked expansion and reorganization of CD4⁺ T-helper populations, including regulatory (T_reg_) cells and multiple clusters characterized by lymphoid follicle-associated features (**Figure 3F**). These T-helper populations exhibit coordinated activation of transcriptional programs associated with follicular helper differentiation and immune orchestration. They are enriched for canonical T_FH_-associated regulators and effector genes, including *Bcl6*, *Cxcr5*, *Icos*, *Il21*, and *Cd40l*. Notably, a subset of these cells, which we termed T_FH_-like cells (**Figure 3F**, red circle), displayed induction of interferon-responsive and inflammatory gene programs (*Ifng*, *Tbx21*, *Cxcr3*), consistent with a functionally activated, T_H1_-skewed phenotype (**Figure 3G**). In addition, these T_FH_-like cells also specifically and strongly express *Tnc*, *Ralyl*, *Sostdc1*, *Cacna1d*, *Tnfsf8*, and *Crppa* (**Figure 3H**). Their expression suggests that this population may have distinct functional properties related to tissue localization, metabolic regulation, or specialized effector functions beyond those of conventional T_FH_ cells. Collectively, this gene signature defines a niche-adapted T_FH_-like CD4⁺ T-cell state that integrates follicular helper identity with activation, interferon responsiveness, and specialized effector signaling, positioning this population to engage and modulate myeloid cells. Importantly, the emergence of this T_FH_-like CD4⁺ T-cell state was largely abolished in *Tet2^-/-^* to *Gsdmd^-/-^* recipients (**Figure 3F**), indicating a requirement for stromal gasdermin D-dependent pyroptosis in its induction and maintenance.

To validate these findings at the protein level, we performed cytokine profiling of bone marrow supernatants in these mice. Consistent with the emergence of T_FH_-like cells, *Tet2^⁻/⁻^* to wild-type recipients showed elevated CD40L, which clustered with other inflammatory mediators, including IL-6, TNF, and IL-1β (**Figure 3I**). This inflammatory cytokine signature was normalized in *Tet2^-/-^* to *Gsdmd^-/-^* recipients, confirming that stromal pyroptosis is required to sustain the T_FH_-associated inflammatory milieu.

### CD40L blockade breaks the feed-forward loop between T_FH_-like cells and Tet2-deficient macrophages

Having identified a stromal pyroptosis-driven myeloid-adaptive immune circuit reinforced by T_FH_-like CD4⁺ T cells expressing CD40L, we next sought to determine whether disruption of this pathway could functionally attenuate Tet2-driven myeloid pathology. Because monocytosis represents an early, robust, and defining feature of Tet2-deficient disease, we reasoned that effective interruption of CD40-CD40L signaling should be reflected by normalization of aberrant monocytosis. To test this, *Tet2^⁻/⁻^* to wild-type chimeras were treated with an anti-CD40L-blocking antibody for three months. CD40L blockade resulted in a significant reduction in circulating monocytes compared with IgG-treated controls, phenocopying the attenuation observed in *Tet2^⁻/⁻^* to *Gsdmd^-/-^* recipients (**Figure 4A**). This was confirmed by flow cytometry, which revealed a significant decrease in the Ly6C^Hi^ inflammatory monocyte subset (**Figures 4B, C**). This inflammatory monocyte population was similarly reduced in the bone marrow and spleen with anti-CD40L treatment (**Figures 4D-F**). The improvement in monocytic composition was accompanied by a significant reduction in splenomegaly (**Figure 4G**). Histopathological examination revealed that anti-CD40L treatment attenuated the hepatic myeloid infiltration, splenic white pulp effacement, and bone marrow hypercellularity characteristic of *Tet2^⁻/⁻^* to wild-type IgG-treated chimeras, restoring tissue architecture comparable to *Tet2^⁻/⁻^* to *Gsdmd^-/-^* recipient (**Figure 4H**). These findings further support CD40-CD40L signaling as a key downstream reinforcement node linking adaptive immune activation to sustained myeloid expansion.

**Figure 4.**
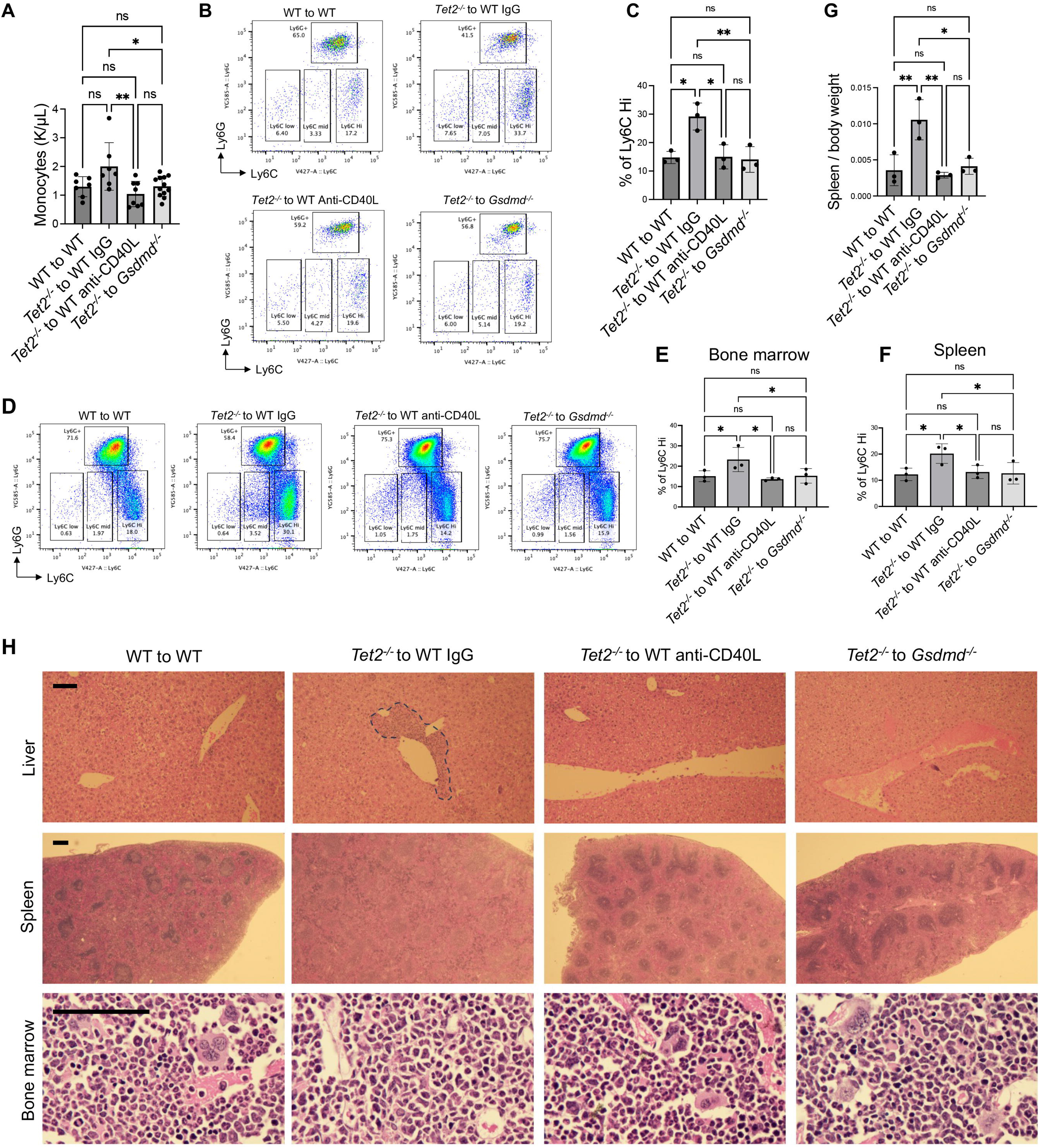
CD40L blockade ameliorates Tet2-deficiency-mediated myeloid proliferation. (A) Peripheral blood monocyte count in *Tet2^⁻/⁻^* to WT chimeras treated with anti-CD40L blocking antibody or IgG control for 3 months. *Tet2^⁻/⁻^* to *Gsdmd^⁻/⁻^* chimeras are shown for comparison. n = 7 in WT to WT group; n = 7 in *Tet2^⁻/⁻^* to WT IgG group; n = 8 in *Tet2^⁻/⁻^* to WT anti-CD40L group; n = 12 in *Tet2^⁻/⁻^* to *Gsdmd^⁻/⁻^* group. (B) Representative flow cytometry plots showing Ly6C^Hi^ inflammatory monocyte frequencies in peripheral blood from indicated treatment groups. (C) Quantification of Ly6C^Hi^ inflammatory monocytes from the indicated groups. n = 3 in each group. (D) Representative flow cytometry plots of Ly6G and Ly6C bone marrow cells in the indicated groups. (E) Quantification of Ly6CHi inflammatory monocyte frequency in bone marrow across the indicated groups. (F) Quantification of Ly6CHi inflammatory monocyte frequency in spleen across different groups. (G) Spleen to body weight ratios in the indicated groups. n = 3 in each group. (H) Histopathological analysis of liver (top row), spleen (middle row), and bone marrow (bottom row) from the indicated chimeras. Tet2⁻/⁻ to WT IgG-treated mice display hepatic myeloid infiltration (dashed outline), splenic white pulp effacement by expanding myeloid elements, and hypercellular bone marrow with increased myeloid-to-erythroid ratio. Scale bars: liver and spleen, 300 μm; bone marrow, 100 μm. All the error bars represent the SEM of the mean. Comparisons among multiple groups were evaluated using a 1-way ANOVA. *p<0.05, **p<0.01. ns: not significant.

To determine whether CD40L blockade alters the immune architecture of Tet2-deficient bone marrow, we next examined global and lineage-resolved single-cell landscapes following anti-CD40L treatment. Unbiased visualization of total bone marrow cells demonstrated that anti-CD40L treatment was associated with a marked reduction in both early-differentiated and mature macrophage populations, accompanied by a parallel decrease in T-cell abundance (**Supplemental Figure 5, Figures 5A, B**). This pattern closely resembled that observed in *Tet2^⁻/⁻^* to *Gsdmd^⁻/⁻^* chimeras. Focused analysis of the macrophage compartment resolved multiple transcriptionally distinct populations spanning early differentiated and mature states (**Figure 5C**). Among these, subpopulation 2 was selectively and prominently expanded in *Tet2^⁻/⁻^* to wild-type IgG-treated chimeras, whereas its abundance was markedly reduced following anti-CD40L treatment and in *Tet2^⁻/⁻^* to *Gsdmd^⁻/⁻^* controls (**Figure 5C**). Transcriptional profiling revealed that macrophage subpopulation 2 was enriched for pathways related to antigen processing and presentation, regulation of immune responses, leukocyte adhesion, and lymphocyte activation (**Figure 5D**). Consistent with this functional annotation, these macrophages upregulated genes encoding the MHC-II presentation machinery (*H2-Aa*, *H2-DMa*, *Cd74*), the immunoproteasome subunit *Psmb10* involved in antigen processing, the co-stimulatory receptor *Cd40*, the inflammatory amplifier *Slamf7*, and the T cell chemoattractant *Ccl5* (**Figure 5E**). This expression profile suggests that these cells are well suited to interact directly with CD4⁺ T cells, consistent with the emergence of T_FH_-like populations within the same microenvironment. Notably, both the expansion of macrophage subpopulation 2 and its associated transcriptional programs were substantially attenuated following CD40L blockade.

**Figure 5.**
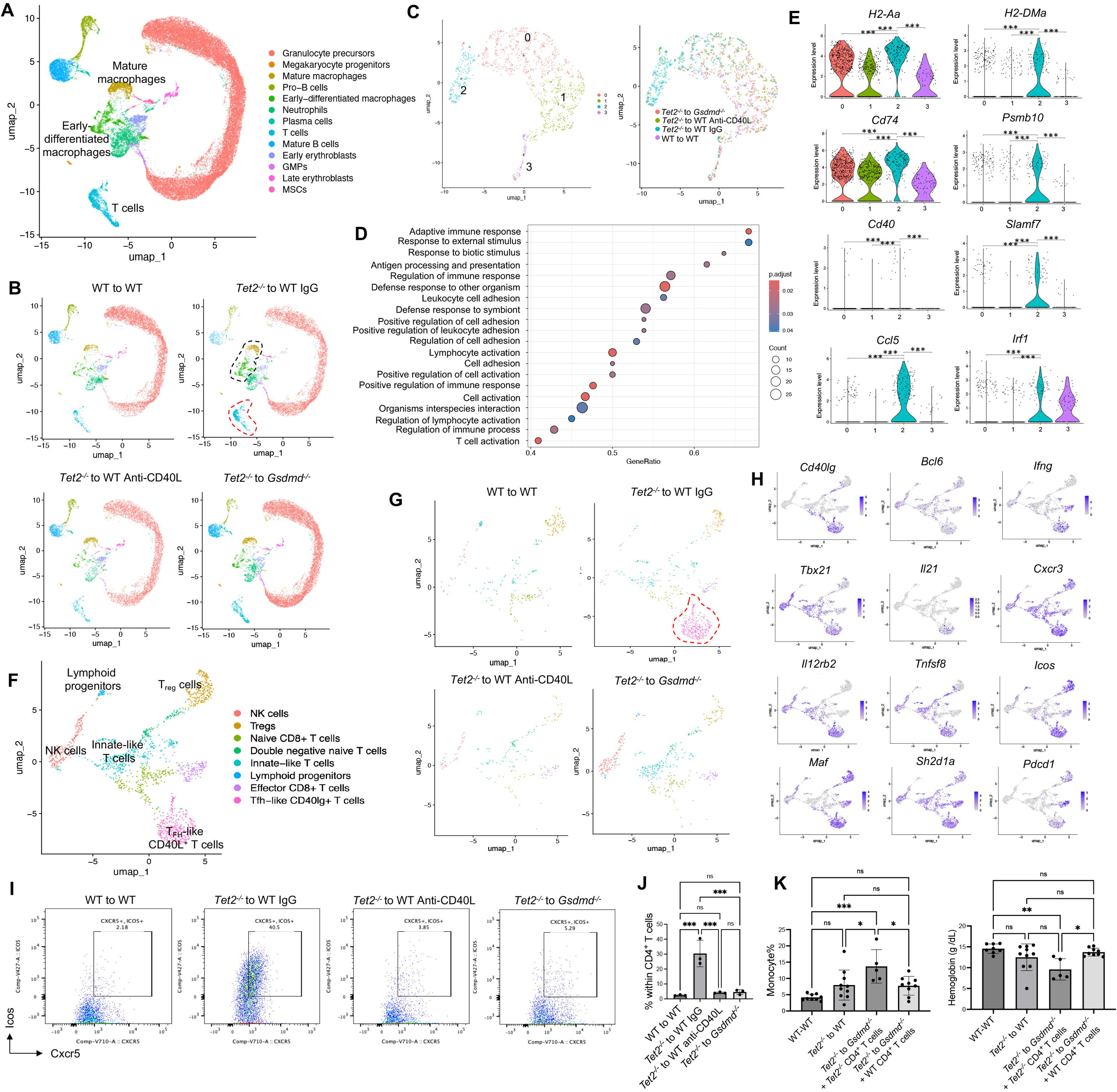
CD40L blockade breaks the feed-forward loop between T_FH_-like cells and Tet2-deficient macrophages. (A) Combined UMAP plot of single-cell RNA sequencing data from total bone marrow cells (post erythroid lysis) from *Tet2^⁻/⁻^* to WT chimeras treated with IgG or anti-CD40L, compared with *Tet2^⁻/⁻^* to *Gsdmd^⁻/⁻^* chimeras. (B) Individual UMAP plot of the indicated group. The black circle indicates macrophage populations; the red circle indicates T-cell populations. (C) UMAP of macrophage subpopulations across indicated groups. (D) Gene Ontology pathway enrichment analysis of genes enriched in macrophage subpopulation 2. (E) Expression of selected genes in macrophage subpopulation 2. (F) Combined UMAP plot of T-cell subpopulations across all groups. (G) Individual UMAP plot of the indicated group. The T_FH_-like population is highlighted in a red circle. (H) Expression of indicated genes across T-cell subpopulations in the indicated groups. (I) Representative flow cytometry plots showing CXCR5⁺ICOS⁺CD4⁺ T cells in bone marrow from the indicated groups. (J) Quantification of CXCR5⁺ICOS⁺CD4⁺ T cells from the indicated groups. n = 3 in each group. (K) Adoptive transfer of CD4⁺ T cells from 8-month diseased *Tet2^⁻/⁻^* mice or age-matched WT controls into *Tet2^⁻/⁻^* to *Gsdmd^⁻/⁻^* chimeras. Monocyte percentage and hemoglobin levels at one-month post-transfer are presented. All the error bars represent the SEM of the mean. Comparisons among multiple groups were evaluated using a 1-way ANOVA. *p<0.05, **p<0.01, and ***p<0.001. ns: not significant.

As expected, analyses of the T cell compartment revealed a distinct T_FH_-like CD40L^+^ population that was selectively expanded in *Tet2^-/-^* to wild-type IgG-treated chimeras. Anti-CD40L treatment markedly reduced this population, restoring the CD4^+^ T cell landscape toward wild-type controls and *Tet2^⁻/⁻^* to *Gsdmd^⁻/⁻^* chimeras (**Supplemental Figure 6, Figures 5F, G, H**). Flow cytometry analyses confirmed a significant reduction of CXCR5^+^ICOS^+^CD4^+^ T cells (**Figures 5I, J**). These data establish that while stromal pyroptosis licenses T_FH_-like cell emergence, CD40-CD40L signaling is required for its sustained maintenance, identifying this axis as the molecular hub of the pathological feedback loop.

Having demonstrated that CD40L blockade attenuates disease by disrupting T cell-macrophage crosstalk, we next asked whether pathogenic CD4⁺ T cells are sufficient to drive disease. We isolated CD4⁺ T cells from *Tet2^⁻/⁻^* mice with established myeloid disease and transferred them into *Tet2^⁻/⁻^* to *Gsdmd^⁻/⁻^* chimeras, which are normally protected from disease despite transplanted with Tet2-deficient HSPCs. Within one month, transfer of disease-derived *Tet2^⁻/⁻^* CD4⁺ T cells restored monocytosis and induced significant anemia compared to wild-type CD4⁺ T cell transfer (**Figure 5K**). The rapid onset and severity of disease likely reflect the acute delivery of preformed pathogenic CD4⁺ T cells. Importantly, *Tet2^⁻/⁻^* to *Gsdmd^⁻/⁻^* chimeras, which also harbor Tet2-deficient T cells, do not spontaneously develop disease, demonstrating that Tet2 loss in T cells is insufficient without prior education by the pyroptotic microenvironment.

### Patients with isolated TET2 mutations exhibit CD4^+^ T-cell skewing and tertiary lymphoid structures

Having defined a stromal pyroptosis-driven, myeloid-adaptive immune axis in murine Tet2-deficient bone marrow, we next asked whether analogous immune remodeling is evident in patients. We reasoned that if TET2-mutant myeloid expansion preferentially licenses CD4⁺ T-cell engagement, this should be reflected in altered CD4:CD8 T-cell composition in patient bone marrow in a mutation-specific manner. To test this prediction, we analyzed CD3⁺ T-cell subsets in bone marrow samples from patients harboring isolated TET2 mutations. These patients predominantly represent early stages of myeloid diseases, including clonal cytopenia of undetermined significance (CCUS) or low-risk myelodysplastic syndromes (**Table 1**), a temporal disease phase in which the effects of a single founding mutation can be examined prior to the frequent acquisition of additional somatic mutations or cytogenetic abnormalities that accompany disease progression. Consistent with our prediction, bone marrow samples from patients with isolated TET2 mutations showed a significant increase in the CD4:CD8 T-cell ratio compared with those of normal bone marrow from lymphoma patients without marrow involvement. In contrast, this skewing was not observed in patients with myeloid disorders harboring isolated DNMT3A mutations, whose CD4:CD8 ratios were comparable to the reference cohort (**Figures 6A, B**). These findings indicate that preferential CD4⁺ T-cell engagement is a feature of early TET2-mutant disease and is not a general property of clonal myeloid disorders.

**Figure 6.**
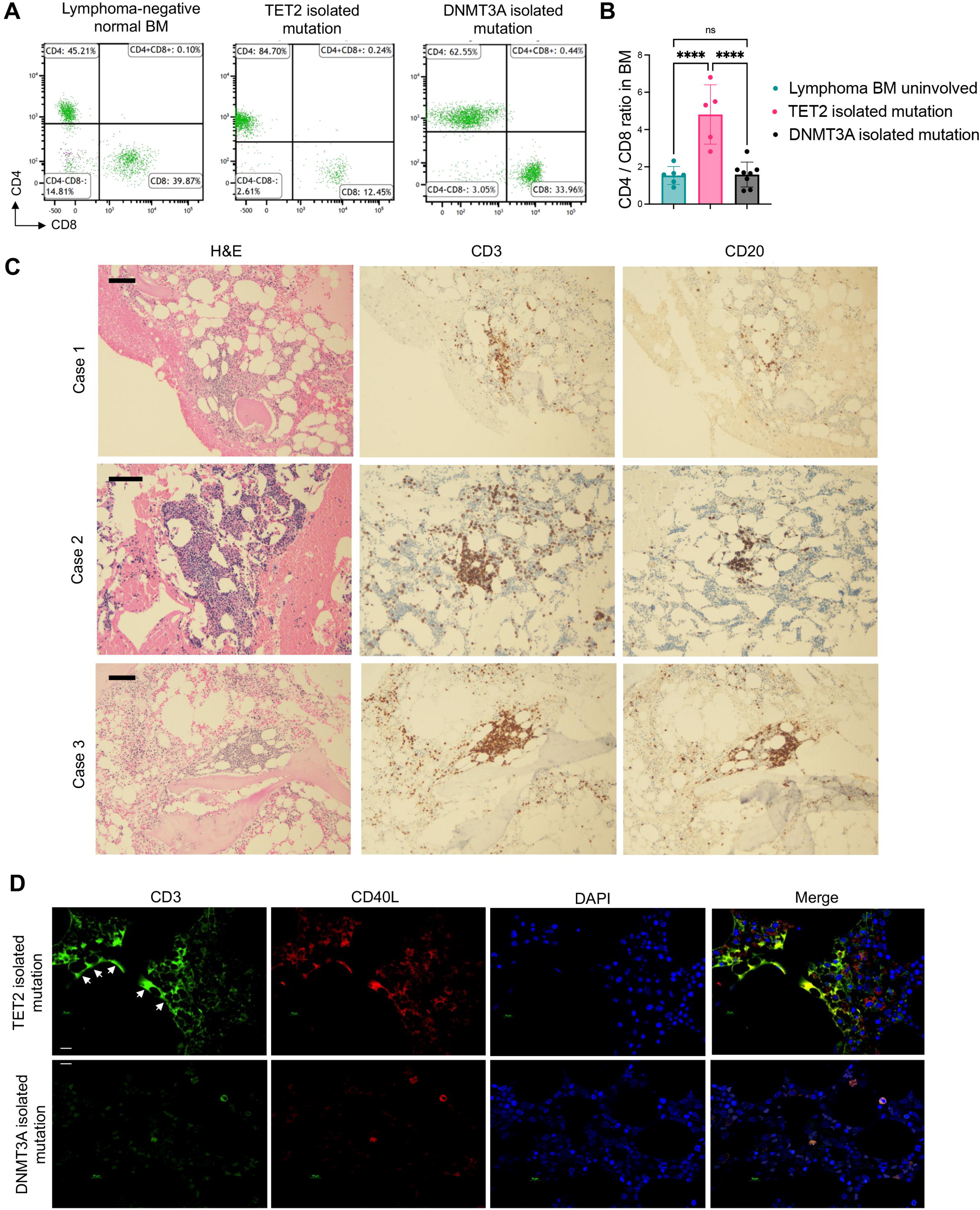
Patients with isolated TET2 mutations exhibit CD4⁺ T-cell skewing and the formation of tertiary lymphoid structures. (A) Representative flow cytometry plots showing CD4⁺ and CD8⁺ T-cell populations in bone marrow from patients with isolated TET2 mutations, isolated DNMT3A mutations, and lymphoma-uninvolved reference controls. (B) Quantification of CD4:CD8 T-cell ratios in bone marrow from the indicated patients. n = 6 in lymphoma-uninvolved group; n = 5 in TET2 isolated mutation group; n = 8 in DNMT3A isolated mutation group. The error bars represent the SEM of the mean. The comparison was evaluated using a 1-way ANOVA. ****p<0.0001. ns: not significant. (C) Representative immunohistochemical staining of bone marrow biopsies showing TLS in patients with isolated TET2 mutations. Serial sections were stained for H&E, CD3 (T cells), and CD20 (B cells). TLS was absent in lymphoma-uninvolved reference samples and isolated DNMT3A-mutant cases analyzed. Scale bars: 150 μm. (D) Representative immunofluorescence staining of bone marrow clot sections from the indicated patients. Sections were stained for CD3 (green), CD40L (red), and DAPI (blue). Merged images are shown on the right. Arrows indicate autofluorescence. Scale bars: 10 μm.

**Table 1:**
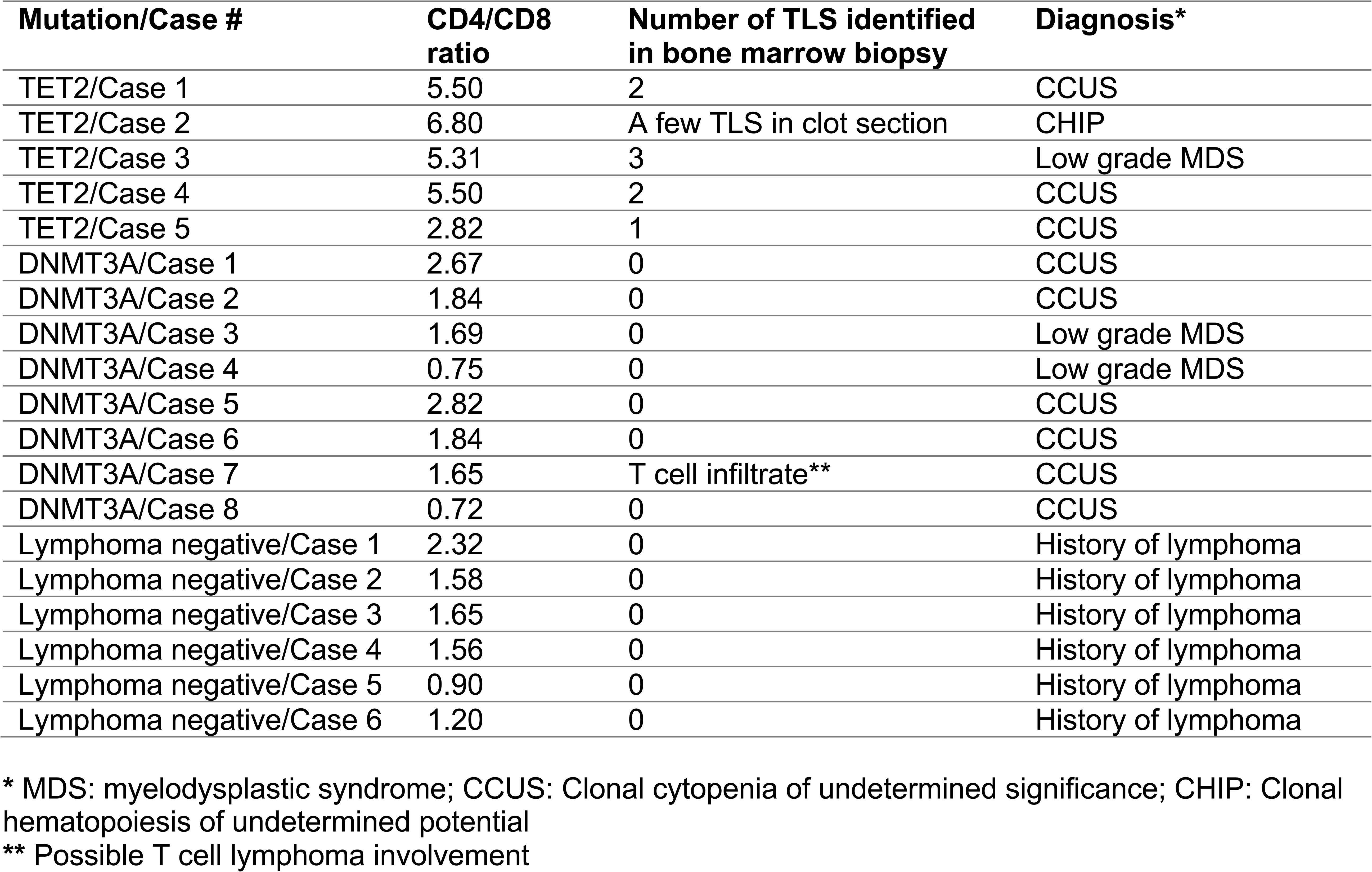
Formation of the tertiary lymphoid structures in TET2-mutated patients.

| **Mutation/Case #** | **CD4/CD8 ratio** | **Number of TLS identified in bone marrow biopsy** | **Diagnosis\*** |
| --- | --- | --- | --- |
| TET2/Case 1 | 5.50 | 2 | CCUS |
| TET2/Case 2 | 6.80 | A few TLS in clot section | CHIP |
| TET2/Case 3 | 5.31 | 3 | Low grade MDS |
| TET2/Case 4 | 5.50 | 2 | CCUS |
| TET2/Case 5 | 2.82 | 1 | CCUS |
| DNMT3A/Case 1 | 2.67 | 0 | CCUS |
| DNMT3A/Case 2 | 1.84 | 0 | CCUS |
| DNMT3A/Case 3 | 1.69 | 0 | Low grade MDS |
| DNMT3A/Case 4 | 0.75 | 0 | Low grade MDS |
| DNMT3A/Case 5 | 2.82 | 0 | CCUS |
| DNMT3A/Case 6 | 1.84 | 0 | CCUS |
| DNMT3A/Case 7 | 1.65 | T cell infiltrate** | CCUS |
| DNMT3A/Case 8 | 0.72 | 0 | CCUS |
| Lymphoma negative/Case 1 | 2.32 | 0 | History of lymphoma |
| Lymphoma negative/Case 2 | 1.58 | 0 | History of lymphoma |
| Lymphoma negative/Case 3 | 1.65 | 0 | History of lymphoma |
| Lymphoma negative/Case 4 | 1.56 | 0 | History of lymphoma |
| Lymphoma negative/Case 5 | 0.90 | 0 | History of lymphoma |
| Lymphoma negative/Case 6 | 1.20 | 0 | History of lymphoma |

\* MDS: myelodysplastic syndrome; CCUS: Clonal cytopenia of undetermined significance; CHIP: Clonal hematopoiesis of undetermined potential
\*\* Possible T cell lymphoma involvement

Strikingly, histopathologic examination of bone marrow biopsies revealed the consistent presence of tertiary lymphoid structures (TLS) in all isolated TET2-mutant cases analyzed. These lymphoid structures were small to medium in size and composed of predominantly small lymphocytes. Immunohistochemical analysis demonstrated prominent enrichment of CD3⁺ T cells within these aggregates, with a lesser contribution of CD20⁺ B cells, indicating a T-cell-dominant lymphoid architecture (**Table 1**, **Figure 6C**). In contrast, TLS was absent in lymphoma-uninvolved reference bone marrow samples and in cases harboring isolated DNMT3A mutations. To determine whether these T cells exhibit features of the T_FH_-like population identified in our murine model, we performed immunofluorescence co-staining for CD3 and CD40L. CD3⁺ T cells within TET2-mutant TLS showed prominent CD40L expression (**Figure 6D**), consistent with the activated, T_FH_-like phenotype observed in Tet2-deficient mice. In contrast, only sparse T cells with CD40L expression were present in DNMT3A-mutant marrow (**Figure 6D**). The consistent detection of T-cell-rich, CD40L-expressing TLS in TET2-mutant bone marrow supports a model in which early TET2-driven disease is associated with localized, spatially organized adaptive immune responses and provides human correlative evidence for the pathogenic role of the CD40-CD40L axis identified in our preclinical studies.

## Discussion

Our study identifies a pyroptosis-dependent stromal-immune axis that links bone marrow niche inflammation to maladaptive remodeling of the adaptive immunity in Tet2-deficient hematopoiesis. We reveal that gasdermin D-mediated pyroptosis exerts distinct mechanisms depending on the underlying genetic lesion. Hematopoietic cell-intrinsic pyroptosis drives MDS pathogenesis in the mDia1/miR-146a-double-deficient model, whereas cell-extrinsic pyroptosis in bone marrow stromal cells is the critical determinant of Tet2-deficient myeloid expansion. This mutation-specific dichotomy demonstrates that inflammatory pathways can be differentially co-opted by distinct myeloid neoplasm-associated genetic abnormalities, with important implications for therapeutic targeting.

Previous studies have shown that TET2-deficient hematopoietic cells are hypersensitive to extrinsic inflammatory signals^16^ and that microbial translocation and IL-6 signaling can promote myeloproliferation in Tet2-deficient hosts^13,31^. Our findings identify pyroptotic stromal cells as a critical cellular source of these inflammatory cues. Tet2-deficient hematopoiesis is inherently biased toward monocytic differentiation^15,28^, and our spatial transcriptomic analyses reveal that these monocytes and macrophages are actively recruited to and cluster in close physical proximity to pyroptotic stromal cells. This spatial juxtaposition positions Tet2-deficient myeloid cells to directly engage with pyroptotic debris. Indeed, we demonstrate that these cells exhibit enhanced phagocytic uptake of pyroptotic stromal material. Importantly, this phagocytosis triggers functional reprogramming toward an activated, MHC-II-high antigen-presenting state, as previously demonstrated^16^, potentially licensing engagement of CD4⁺ T cells and bridging innate inflammation to adaptive immune remodeling. Thus, the monocytic bias of Tet2-deficient hematopoiesis, combined with enhanced phagocytic capacity and antigen presentation, creates a cellular substrate uniquely positioned to bridge stromal pyroptosis to adaptive immune activation, a mechanism that may distinguish progressive from indolent TET2-mutant clones. This supports a “living with the mutation” paradigm: patients with TET2-mutant hematopoietic cells can indefinitely live without clinical consequences if the niche remains non-pyroptotic. Therapeutically, this suggests that targeting niche inflammation or the maladaptive immunity could stabilize the disease without requiring the eradication of the mutant clone.

A key finding of this study is the emergence of a T_FH_-like CD4⁺ T-cell population in the pyroptotic bone marrow niche. These cells express not only canonical T_FH_ markers but also interferon-responsive and tissue-interaction programs, distinguishing them from conventional germinal center T_FH_ cells. This hybrid transcriptional state suggests adaptation to the inflammatory bone marrow microenvironment rather than classical germinal center reactions. The dependence of this population on stromal gasdermin D indicates that pyroptosis-derived signals, rather than Tet2 deficiency itself, are required for their emergence and maintenance. Importantly, the T_FH_-like population was not merely a bystander but functionally contributed to disease pathogenesis, as CD40L blockade attenuated both T_FH_-like cell expansion and inflammatory monocytosis.

We found that the CD40-CD40L axis emerges as a central node linking adaptive immunity to Tet2-deficient myeloid expansion. CD40L from T_FH_-like cells engages CD40 on Tet2-deficient macrophages, creating a self-amplifying circuit that can be disrupted to attenuate disease. The translational relevance of these findings is supported by our observation that patients with isolated TET2 mutations exhibit CD4:CD8 T-cell skewing and T-cell-rich tertiary lymphoid structures, features absent in DNMT3A-mutant cases, suggesting mutation-specific immune signatures with potential diagnostic utility. Therapeutically, our data provide rationale for targeting T-cell-macrophage crosstalk via CD40L blockade, agents already in clinical development for inflammatory and autoimmune diseases^35–37^. The mutation-specific nature of this stromal-immune axis underscores the importance of genotype-directed therapeutic strategies in myeloid neoplasms.

Several questions remain for future investigation. The precise signals released by pyroptotic stromal cells that recruit and activate Tet2-deficient macrophages remain to be fully characterized. While we demonstrate spatial proximity and functional interaction, the specific DAMPs, alarmins, or cytokines mediating this crosstalk remain to be defined. Additionally, the antigens presented by MHC-II-high Tet2-deficient macrophages to T_FH_-like cells are unknown; whether these represent self-antigens, neoantigens, or pyroptosis-derived material warrants further investigation. Finally, whether the T_FH_-like population identified here represents a stable lineage or a transient activation state, and whether it has counterparts in other inflammatory bone marrow conditions, remains to be determined.

## Methods

### Animals

All mouse experiments were performed in accordance with protocols approved by the Northwestern University Institutional Animal Care and Use Committee (IACUC; protocol IS00028219). *Gsdmd*^-/-^ mice were obtained from Feng Shao (National Institute of Biological Sciences, Beijing). *Tet2* floxed mice (#017573), *Vav-iCre* mice (#008610), and *miR-146a* knockout mice (#016239) were purchased from the Jackson Laboratory. To generate hematopoietic-specific *Tet2* knockout mice, *Vav-iCre* mice were crossed with *Tet2* floxed mice to obtain *Vav-iCre*^+^;*Tet2^fl/fl^*animals (hereafter referred to as *Tet2*^-/-^). mDia1/miR-146a double-knockout (DKO) mice were generated as described previously^7,10,11^. Congenic CD45.1 mice were obtained from Charles River. Mice were maintained in a standard institutional holding facility under routine housing conditions. For transplantation-based studies, donor and recipient mice were age-matched, with cohorts comprising approximately equal numbers of male and female animals. Genotyping was performed by PCR using genomic DNA isolated from tail biopsies. Primer sets and genotyping strategies followed the recommendations provided by the Jackson Laboratory for the relevant strains, or as described previously^7,10,11^.

### Human samples collection

Human bone marrow aspirate samples, clot section and core biopsy blocks were obtained from leftover diagnostic specimens at Northwestern Memorial Hospital. The study protocol was approved by Northwestern University’s Institutional Review Board (IRB ID: STU00217116). Human sample embedding and sectioning were performed at Northwestern University’s Pathology Core Facility.

### Bone Marrow Transplantation

Recipient mice were 10-14 weeks of age at the time of transplantation. Both sexes were used as indicated in individual experiments, with animals randomly assigned to experimental groups. Recipient mice were conditioned with a single dose of 10 Gy total body irradiation using a cesium gamma irradiator. Donor cells were transplanted within 12-24 hours following irradiation.

Donor hematopoietic cells were isolated from the bone marrow of long bones (femurs and tibias). Lineage-negative (Lin⁻) hematopoietic stem and progenitor cells were enriched using the Mouse Lineage Cell Depletion Kit (Miltenyi Biotec, 130-110-470) according to the manufacturer’s instructions. Enriched Lin⁻ cells were resuspended in sterile phosphate-buffered saline (PBS), and approximately 2 × 10^5^ Lin⁻ cells were transplanted per recipient mouse via retro-orbital injection in a total volume of 80-100 μL. To reduce donor-to-donor variability, bone marrow cells from three donors were pooled prior to enrichment and transplantation.

To reduce the risk of infection following irradiation, mice received sulfamethoxazole-containing antibiotic water (0.8 mg/mL), beginning 1 day prior to irradiation and continuing through day 21 post-transplantation, after which animals were returned to standard autoclaved drinking water. Animals were monitored daily for signs of distress and provided supportive care as needed.

Peripheral blood was collected by tail snip into capillary tubes and immediately transferred into EDTA-coated collection tubes (MiniCollect; Greiner Bio-One). Complete blood counts were obtained using a Heska Element HT5 hematology analyzer. For survival analyses, animals were euthanized upon reaching a predefined moribund stage to minimize suffering. Moribund stage was defined as the presence of one or more of the following clinical signs: severe lethargy or unresponsiveness to external stimuli; inability to ambulate or maintain upright posture; failure to access food or water; marked weight loss (>20% of baseline body weight); labored or irregular respiration; severe dehydration; or persistent hypothermia. Mice meeting these criteria were humanely euthanized, and the date of euthanasia was recorded as the survival endpoint.

### Flow Cytometry

Flow cytometry assays were performed as previously described^11,38,39^. Briefly, single-cell suspensions were prepared from peripheral blood (PB), bone marrow (BM), and spleen. For PB and BM, red blood cells were removed using Red Cell Lysis Buffer (eBioscience, 00-4300-54), and remaining cells were passed through a 40 μm cell strainer to remove aggregates. Spleens were mechanically dissociated by mincing and homogenization using the frosted ends of microscope slides in MACS buffer, followed by filtration through a 40 μm cell strainer.

For analyses of surface markers, freshly prepared cells were stained and analyzed without fixation. For intranuclear markers, cells were first stained with antibodies targeting cell-surface markers, then fixed and permeabilized with the Foxp3/Transcription Factor Staining Buffer Set (eBioscience, 00-5523-00), followed by intranuclear staining according to the manufacturer’s instructions.

Data were acquired on a BD Symphony S5 SE and analyzed using FlowJo v10.0. Compensation controls were prepared using compensation beads (UltraComp eBeads, eBioscience, #01-3333-41) stained with the same antibodies used in each assay, and spectral unmixing was performed in FlowJo using the built-in spectral unmixing algorithm. Antibody panels (including clones, fluorophores, and vendors) are provided in the **Supplemental Table 1**.

### Immunoblotting

Total bone marrow cells were lysed in RIPA buffer supplemented with phosphatase inhibitor (PhosSTOP; Roche). Protein concentration was determined by BCA assay. Lysates were reduced with 2-mercaptoethanol and resolved by SDS–PAGE using 4-12% gradient gels (Bio-Rad), followed by wet transfer to a membrane. Membranes were probed with an anti-N-terminal GSDMD antibody (ab219800; Abcam) and HSC70 (Santa Cruz, #sc-7298) as a loading control. Immunoreactive bands were detected by enhanced chemiluminescence (ECL) and visualized using a Bio-Rad ChemiDoc XRS imaging system.

### Hematoxylin and eosin (H&E) stain

Mouse sternum, spleen, liver, and lung were harvested and fixed in 10% neutral-buffered formalin overnight. Fixed tissues were paraffin-embedded, sectioned, and processed for H&E staining by the Mouse Histology and Phenotyping Laboratory at Northwestern University.

### Immunofluorescence

Femurs were fixed in 4% paraformaldehyde at room temperature for 72 hours, embedded in 8% gelatin (8 g gelatin in PBS), and stored at −80°C for 24 hours prior to cryosectioning, as described previously^11,40^. Cryosectioning was performed at −25°C to generate 100 μm-thick cross sections, which were collected onto microscope slides and allowed to equilibrate to room temperature. During equilibration, differential thermal expansion between bone and marrow separates the bone from the bone marrow while preserving marrow structural integrity.

Sections were permeabilized in 0.25% Triton X-100 and blocked in 5% goat serum. Sections were incubated with primary antibodies at room temperature for 1 hour, then overnight at 4°C. Primary antibodies included rabbit anti-Gsdmd (Invitrogen, PA5-115330) and mouse anti-CD271 (Invitrogen, 14-9400-82). Sections were then incubated with goat anti-mouse IgG (H+L) cross-adsorbed secondary antibody (Alexa Fluor 488) and goat anti-rabbit IgG (H+L) cross-adsorbed secondary antibody (Alexa Fluor 647). Endothelial structures were labeled with biotin-UEA1 (Vector Laboratories, B0-1065-2) and streptavidin-alexa Fluor 568. Sections were optically cleared using a fructose-based clearing method and mounted as described previously^41^. Imaging was performed on a Nikon A1R laser-scanning confocal system using a 20× water-immersion objective (CFI Plan Apochromat Lambda D). Antibody panels (including clones, fluorophores, and vendors) are summarized in the **Supplemental Table 2**.

### Single-cell RNA sequencing

Single-cell suspensions for scRNA-seq were prepared from mouse bone marrow (BM) by flushing femurs and tibias, followed by red blood cell lysis using RBC Lysis Buffer (eBioscience). For MSC-enriched scRNA-seq, bones were minced and digested with collagenase type II (STEMCELL Technologies) at 37°C for 1 hour, and the resulting cell suspension was passed through a 40 μm cell strainer to remove debris and aggregates. Single-cell libraries were generated using the 10x Genomics Chromium platform, targeting ∼100,000 cells per sample. Sequencing data were processed using Cell Ranger using the GRCm39 reference genome, and downstream analyses were performed using Seurat.

### Mouse bone marrow cytokine analysis

Bone marrow fluid was harvested from a single femur by centrifugation into 500 μl PBS. Samples were centrifuged to remove cellular debris, and the supernatant was transferred to ultra-low-binding tubes (Eppendorf, EP022431081) and stored at −80 °C until analysis.

Cytokines, chemokines, and other soluble factors in mouse bone marrow fluid were measured by Eve Technologies using bead-based multiplex immunoassays (Discovery Assay® platform). Samples were analyzed using the Mouse Cytokine/Chemokine 44-Plex Discovery Assay® Array (MD44), Mouse High Sensitivity T Cell 18-Plex Discovery Assay® Array (MDHSTC18), and Mouse Cardiovascular Disease Panel 2 9-Plex Assay (MCVD2-09-107). Bone marrow fluid was analyzed without dilution.

### Cryosection of the mouse femur without decalcification

This procedure was performed to prevent decalcification-induced RNA degradation in spatial transcriptomic studies. The femur was fixed in 4% paraformaldehyde at room temperature for 72 hours, followed by embedding in 8% gelatin (8 g gelatin dissolved in PBS) and storage at −80°C for 24 hours prior to cryosectioning. Cryosectioning was performed at −25 °C, and the femur was sectioned into 100 µm-thick slices that were mounted onto coated microscope slides. The slides were subsequently equilibrated to room temperature. During this equilibration process, differences in the coefficients of thermal expansion between bone and bone marrow facilitated the mechanical separation of bone from marrow while preserving the structural integrity of the bone marrow. The isolated bone marrow was then embedded in paraffin and sectioned onto slides specifically designed for spatial transcriptomic analysis.

### Xenium spatial transcriptomics

Xenium spatial transcriptomics reagents and instrumentation were obtained from 10x Genomics (Pleasanton, CA, USA). Paraffin-embedded tissue sections (5 µm) were mounted onto Xenium capture areas and incubated at 42°C for 3 hours, then dried overnight. Slides were subsequently stored at room temperature in a desiccation chamber until deparaffinization.

Slide processing and Xenium Analyzer loading were performed using the Xenium Slide & Sample Preparation Reagent Kit (PN1000460), Xenium Decoding Consumables (PN1000487), and Xenium Decoding Reagent (PN1000461). Sections were deparaffinized and de-crosslinked according to the manufacturer’s protocol (CG000580). Custom probe panels targeting preselected genes were hybridized to tissue sections overnight, followed by probe ligation and signal amplification according to the manufacturer’s instructions (CG000582). Autofluorescence quenching and nuclear staining were performed prior to imaging and signal decoding on the Xenium Analyzer. All Xenium slide processing and analyzer loading were conducted at the NUSeq Core Facility at Northwestern University.

### Workflow of Xenium Spatial Gene Expression

Xenium data were processed using Space Ranger (10x Genomics; https://www.10xgenomics.com/support/software/space-ranger/latest/tutorials/count-ffpe-tutorial) to generate FASTQ-derived outputs. Cell feature matrices, cell and nuclear boundary files, transcript coordinates, and spatial morphology images were exported for downstream analysis using Giotto Suite (v4.0.5) in R (v4.3.3). For the Xenium analysis, a Giotto Xenium object was created by loading the probe, expression matrix, and image files, followed by integrating metadata and cell and nuclear boundary information. Quality filtering was performed using the parameters: expression threshold = 1, minimum detected features per cell = 5, and features detected in ≥3 cells. Data were then normalized with a scale factor of 5,000. Feature and cell-level statistics were calculated prior to dimensionality reduction by identifying highly variable features and performing principal component analysis (PCA).

Two-dimensional embeddings were generated using t-distributed stochastic neighbor embedding (t-SNE) and uniform manifold approximation and projection (UMAP) based on the first 10 principal components. A nearest-neighbor graph was constructed, and cells were clustered using the Leiden community detection algorithm at resolutions ranging from 0.2 to 0.5. Spatial mapping of clusters was performed to visualize in situ distributions. Cluster marker genes were identified using the scran package.

Cell type annotation was performed by integrating marker gene signatures, reference datasets from the Mouse Cell Atlas, and metadata from matched single-cell RNA-seq datasets. Annotated cell types were mapped back to spatial coordinates to visualize their in situ distributions. A spatial Delaunay network (maximum distance = 400) was constructed to model cell-to-cell spatial relationships. Cell proximity enrichment analysis with 1,000 simulations was performed to infer potential cell–cell interactions. All analysis scripts and detailed parameter settings are available at the project repository (https://github.com/kehanren-sudo/Xenium-Analysis).

### In vivo anti-CD40L blockade

For in vivo CD40L blockade, mice were treated with anti-mouse CD40L antibody (clone MR-1, Bio X Cell, BE0017-1) at a dose of 8 μg/g body weight by intraperitoneal injection twice weekly. Treatment was initiated 1 month after bone marrow transplantation and continued for 8 weeks. Control animals received Armenian hamster IgG isotype control antibody (Bio X Cell, BE0091) on the same schedule. Experimental endpoints were collected at 3 months post-transplantation.

### Macrophage-MSC engulfment assay

Bone marrow-derived macrophages (BMDMs) were generated from 3-month-old mice of the indicated genotypic background using a previously described method^42^. Mouse mesenchymal stromal cells (MSCs) were enriched from the bone marrow of 3-month-old mice of the indicated genotypic background using the mouse MesenCult™ Expansion Kit (STEMCELL Technologies, #05513). BMDMs were seeded in 12-well plates at a density of 1 × 10^5^ cells per well and allowed to adhere overnight. MSCs were labeled while adherent with CellTrace Far Red (Invitrogen, C34564) according to the manufacturer’s instructions. Labeled MSCs were then treated overnight with recombinant mouse S100A9 protein (R&D Systems, 2065-S9; 300 ng/mL) and uric acid (MilliporeSigma, U0881-10G; 150 μM) under hypoxic culture conditions. The following day, MSCs were treated with nigericin (5 μg/mL) for 15 minutes, detached, and resuspended in fresh MesenCult Expansion Basal Medium without MesenPure Supplements. Cell viability under these conditions remained >90%.

Treated MSCs were counted and added to BMDM cultures at a 1:1 ratio under normoxic conditions. After overnight co-culture in MesenCult Expansion Basal Medium, cells were harvested using ReLeSR (STEMCELL Technologies, #100-0483) in combination with physical detachment, stained with the indicated antibodies, and analyzed by flow cytometry to assess engulfment of CellTrace-labeled MSCs by macrophages.

### CD4^+^ T cell transfer assay

Adoptive CD4 T-cell transfer experiments were performed in *Tet2^-/-^* to *Gsdmd^-/-^* BMT mice at 2 and 4 weeks post-transplantation. Donor CD4^+^ T cells were isolated from the bone marrow of 8-9-month-old *Vav-iCre;Tet2^f/f^* mice or age-matched *Tet2^f/f^*littermate control mice. Bone marrow cells were harvested from donor mice and processed into single-cell suspensions prior to CD4^+^ T-cell purification. CD4^+^ T cells were purified using the MACS CD4^+^ T Cell Isolation Kit for mouse (Miltenyi Biotec, 130-104-454) according to the manufacturer’s instructions.

For each transfer assay, donor CD4^+^ T cells were pooled from at least five *Vav-iCre;Tet2^f/f^* mice or five *Tet2^f/f^* control mice. Purified cells were resuspended in sterile PBS, and 5 × 10^5^ CD4^+^ T cells in a total volume of 80 μL were administered to each recipient by retro-orbital injection at each time point. Transfers were performed at 2 and 4 weeks after BMT, resulting in a cumulative total of 1 × 10^6^ CD4^+^ T cells transferred per recipient mouse.

## Data availability

The single-cell RNA sequencing data generated in this study have been deposited in GEO under accession number GSE324446. Spatial transcriptomic analysis scripts and detailed parameter settings are available at the project repository (https://github.com/kehanren-sudo/Xenium-Analysis).

## Acknowledgments

We thank the Mouse Histology and Phenotyping Laboratory, the Pathology Core Facility, the Flow Cytometry Facility, the Center for Advanced Microscopy, and the NUSeq Core Facility at Northwestern Feinberg School of Medicine for their assistance with various tissue preparations and assays. We also thank Feng Shao at the National Institute of Biological Sciences, Beijing, for providing gasdermin D knockout mice. This work was supported by the National Heart, Lung, and Blood Institute (NHLBI) grant R01-HL169507 (P.J.) and National Institute of Diabetes and Digestive and Kidney Disease (NIDDK) grant R01-DK138205 (P.J.). K.R. is a recipient of the NIH K99 pathway to independent award (K99CA289959) and the EvansMDS foundation Young Investigator award. H.B. is a recipient of the F32 Ruth L. Kirschstein Postdoctoral Individual National Research Service Award (F32-HL170648).

## Author contributions

K.R., X.H., E.L., P.W., H.B., I.A., L.S., C.M.W., H.N., and J.Y. performed the experiments and interpreted data. C.M.W. performed the sequencing experiments. Y.L., B.V., M.S., and P.J. collected and interpreted human specimens and data. K.R. and X.H. analyzed the sequencing data. K.R., X.H., and P.J. designed the experiments, interpreted data, and edited the manuscript.

D.F. and W.C. interpreted the data and edited the manuscript. P.J. wrote the manuscript.

## Ethics declarations

The authors declare no relevant competing interests related to this work.

## Notes

### Competing Interest Statement

The authors have declared no competing interest.

