## Supplemental figures and tables for "Stromal Gasdermin D-mediated Pyroptosis Drives Maladaptive CD4⁺ T-cell Remodeling in Tet2-Deficient Hematopoiesis"

### Supplementary Data

### Figures and Figure Legends

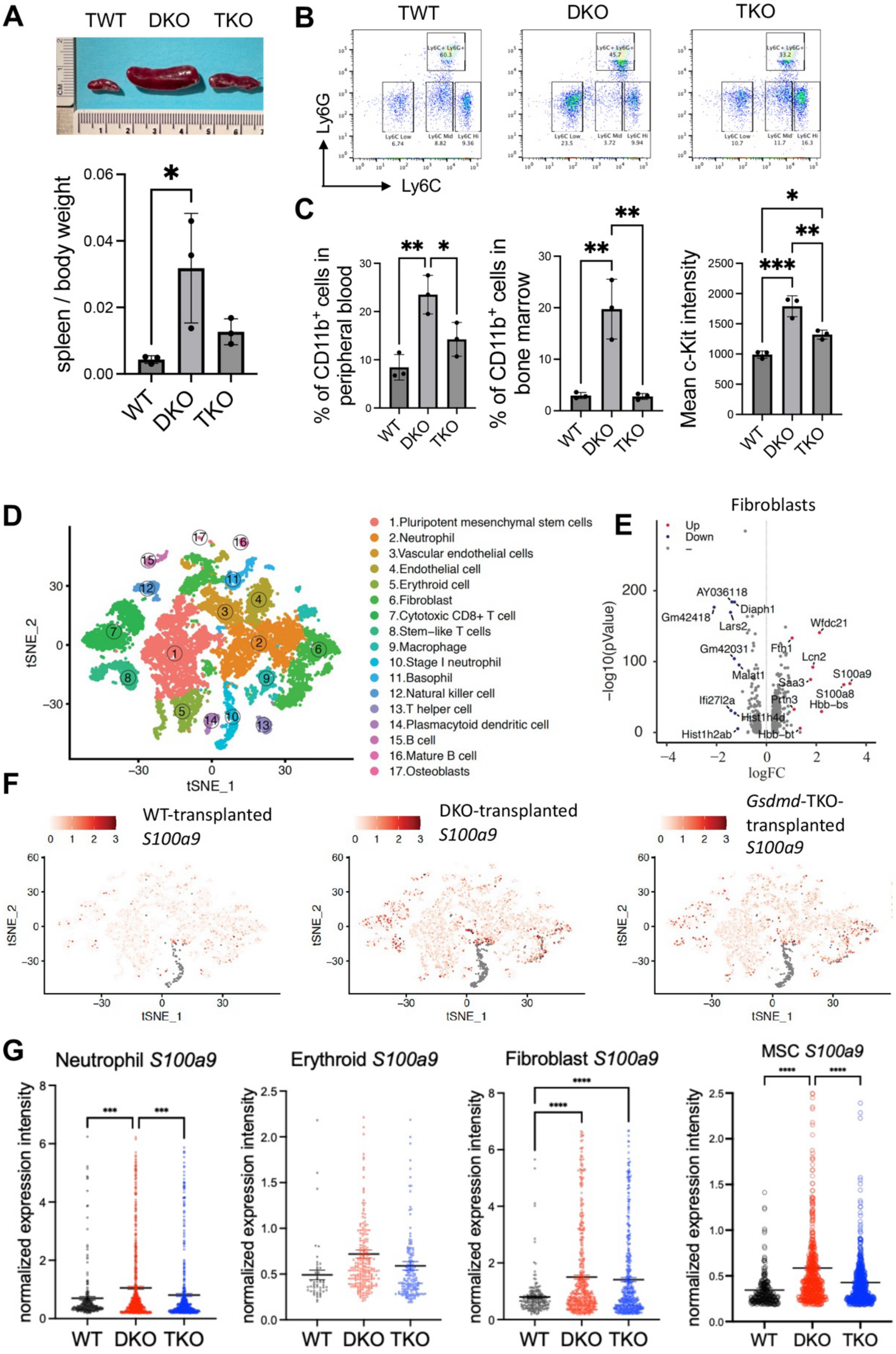

**Supplemental Figure 1. Gasdermin D is important in remodeling bone marrow stromal cells in the DKO model.** (A) Representative pictures of the spleens of the recipient mice at 9 months post transplantation of the indicated donor bone marrow HSPCs. The spleen-to-body weight ratio of the indicated mice is shown below. N = 3 in each group. TWT: triple wild type; DKO: mDia1/miR-146a double knockout; TKO: DKO + *Gsdmd* knockout. (B) Representative flow cytometry plots of Ly6G and Ly6C in the peripheral blood of the indicated mice in A. (C) The percentage of Ly6G$^{-}$ Ly6C$^{low}$ immature monocytic population in CD11b$^{+}$ cells in the peripheral blood and bone marrow of the recipient mice 9 months post transplantation of the indicated HSPCs. Their c-Kit levels are shown on the right. n = 3 in each group. (D) Merged tSNE plots of the partial lineage-depleted bone marrow stromal cells from mice in C. Clusters of annotated cell populations are illustrated. (E) Volcano plot of differentially expressed genes comparing fibroblasts from recipient mice transplanted with wild-type or DKO total bone marrow cells. Up and down-regulated genes in DKO transplanted mice are illustrated. (F) S100a9 expression profiles across cell populations in partially lineage-depleted stromal cells from mice in C. (G) Statistical analyses of S100a9 expression in the indicated cell populations from the indicated mice in F. Cells with detectable S100a9 expression were included in the analyses. All the error bars represent the SEM of the mean. Comparisons among multiple groups were evaluated using a 1-way ANOVA. *p<0.05, **p<0.01, ***p<0.001, and ****p<0.0001.

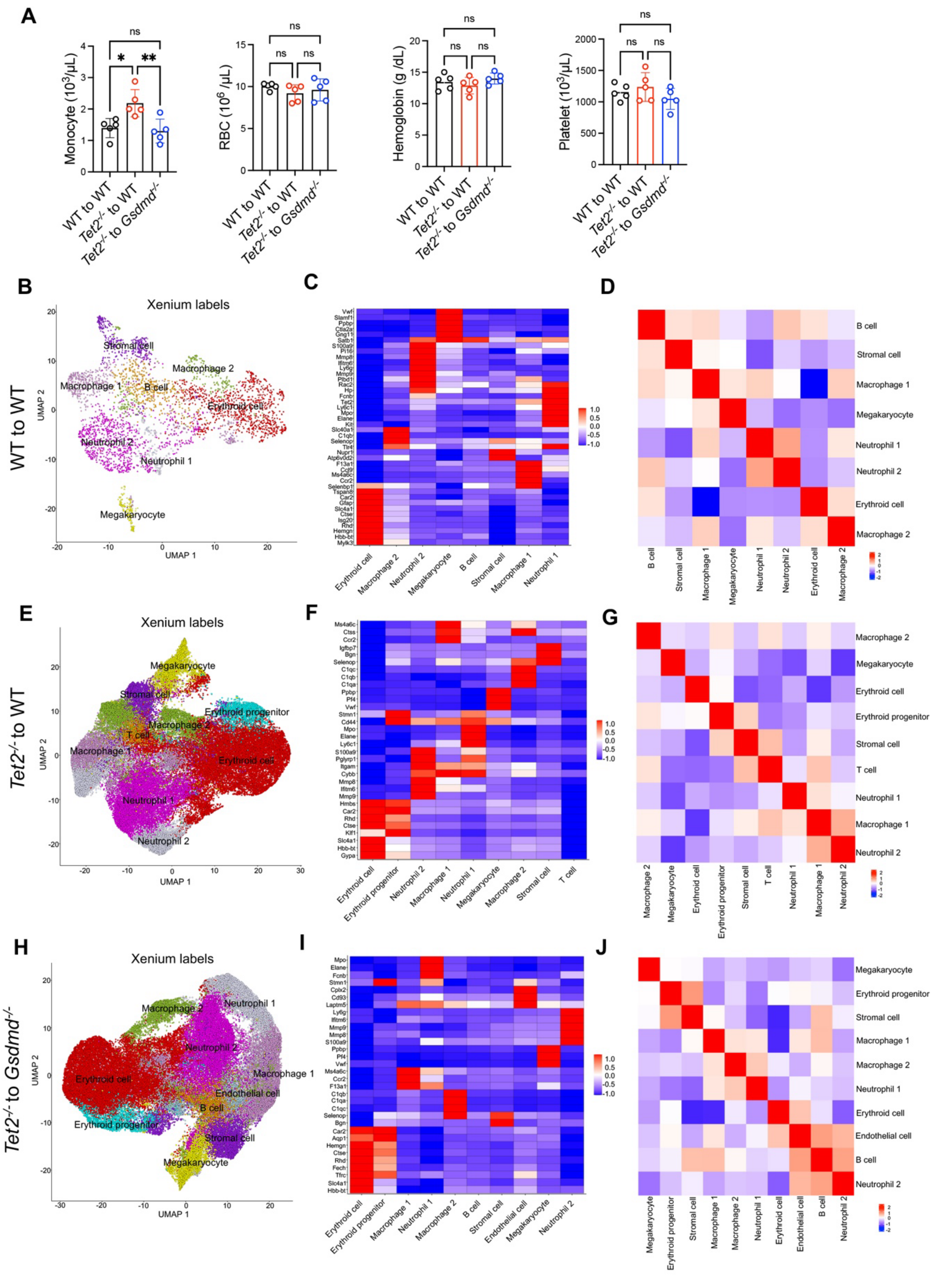


**Supplemental Figure 2. Stromal pyroptosis in Tet2-deficient myeloid expansion.** (A) Monocyte count, RBC count, hemoglobin, and platelet count of the indicated mice at 3 months post-transplant. n = 5 in each group. (B) UMAP plot derived from Xenium subcellular spatial transcriptomic data showing different cell types in the bone marrow of the indicated recipient mice. (C) Specific marker genes of all cell types in B. (D) A heatmap showing interactions between two cell types from the Xenium data in C. (E-G) Same as B-D, except that different indicated recipient mice were illustrated. (H-J) Same as B-D, except that different indicated recipient mice were illustrated.

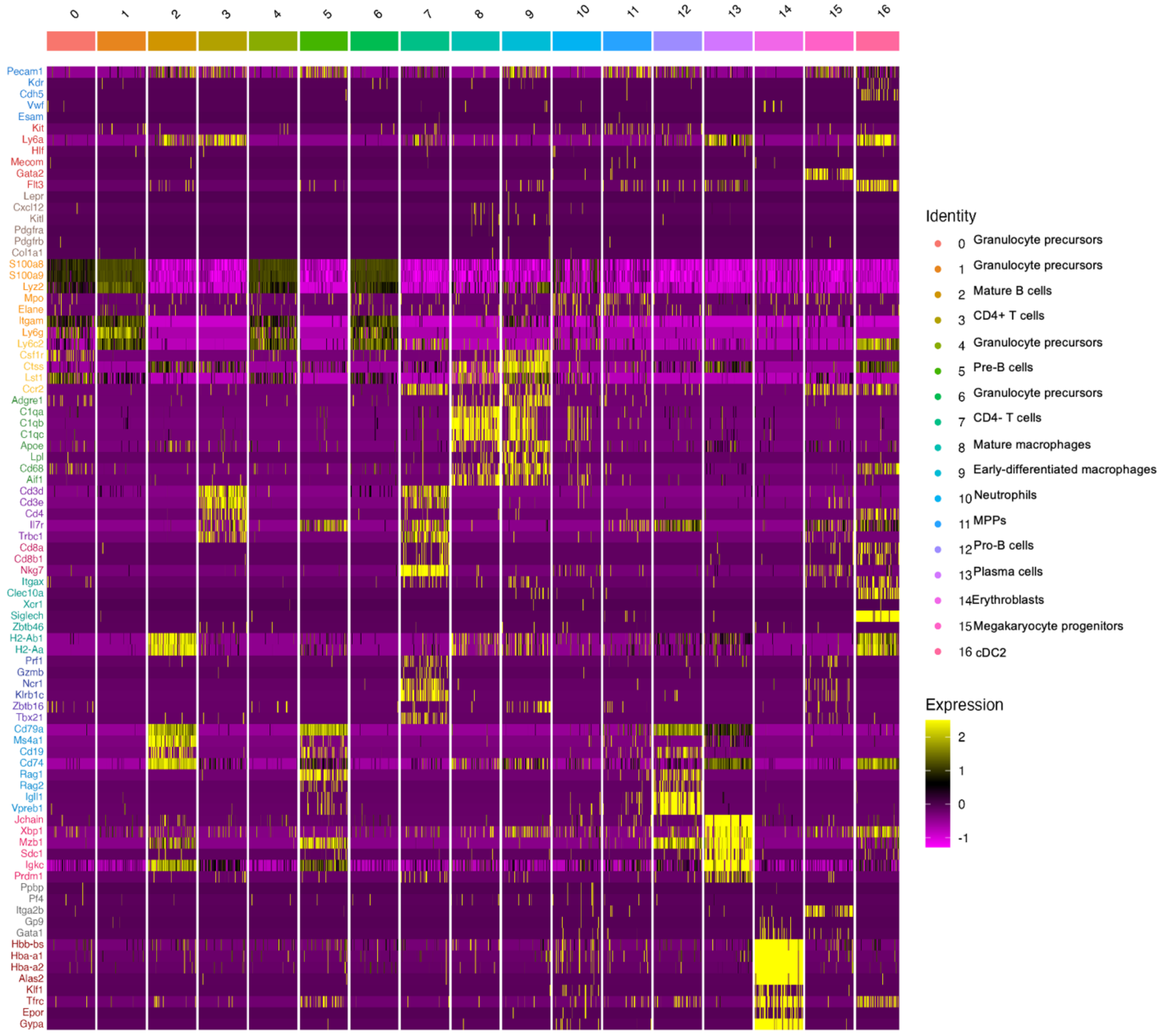


**Supplemental Figure 3. Marker gene expression of the indicated cell types.** Data from combined single-cell RNA sequencing from total bone marrow cells (after erythroid lysis) from WT to WT, *Tet2*$^{-/-}$ to WT, and *Tet2*$^{-/-}$ to *Gsdmd*$^{-/-}$ recipients at 9 months post-transplant.

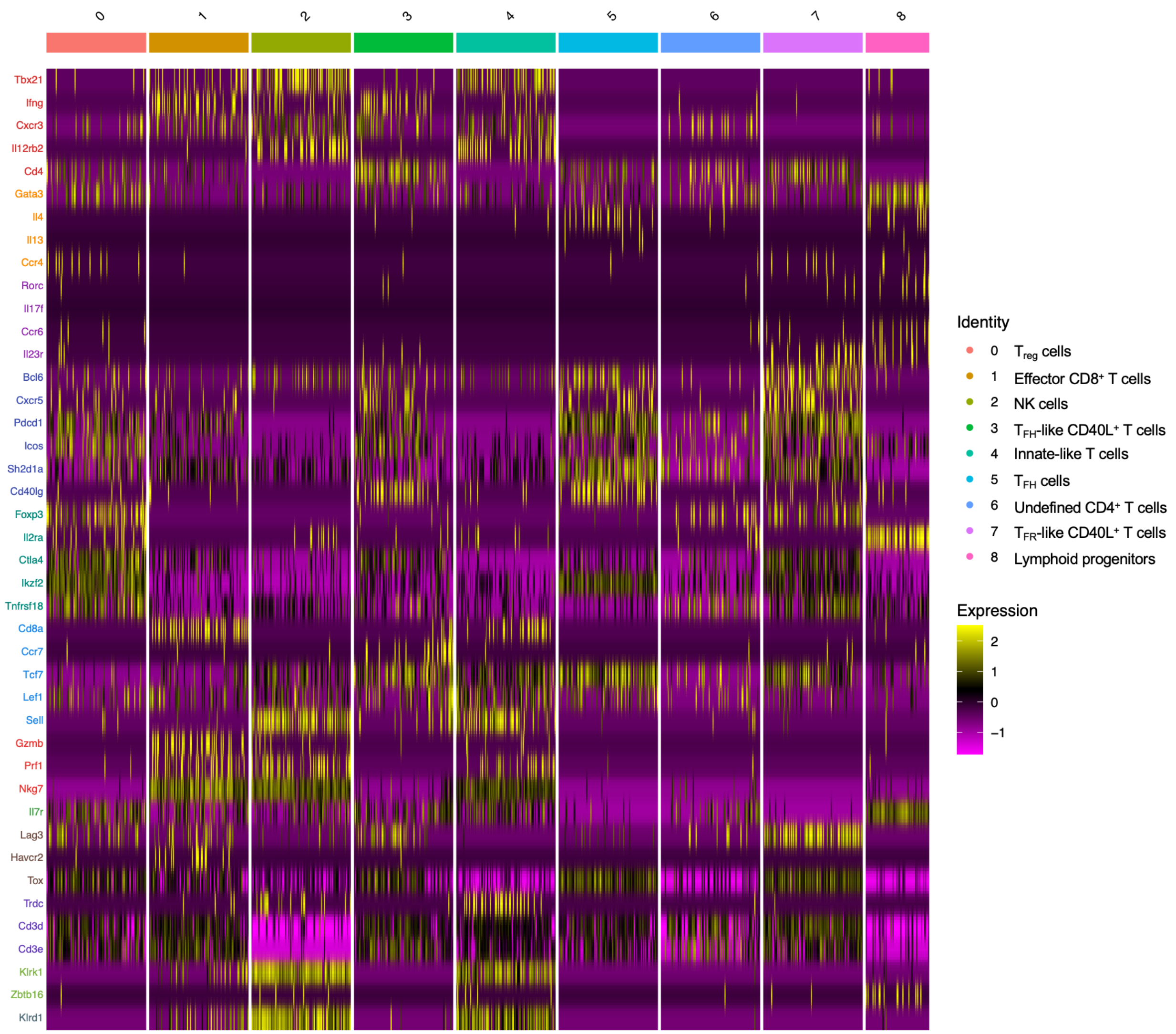


**Supplemental Figure 4. Marker gene expression of indicated T cell subtypes.** Data from further analyses of T-cell subpopulations of supplemental Figure 3.

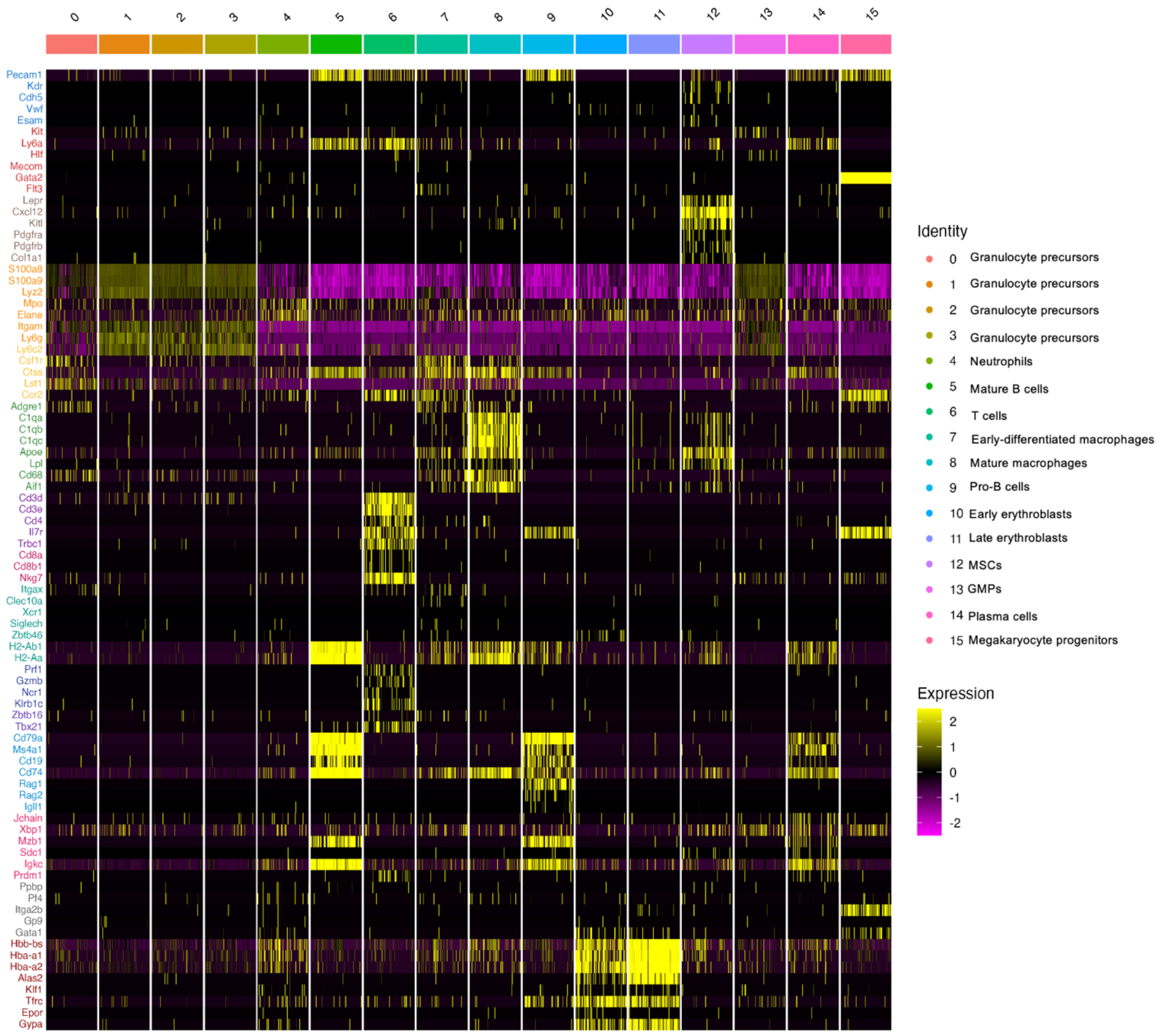


**Supplemental Figure 5. Marker gene expression of indicated cell types.** Data from combined single-cell RNA sequencing from total bone marrow cells (post erythroid lysis) from *Tet2*$^{-/-}$ to WT chimeras treated with IgG or anti-CD40L, compared with *Tet2*$^{-/-}$ to *Gsdmd*$^{-/-}$ chimeras.

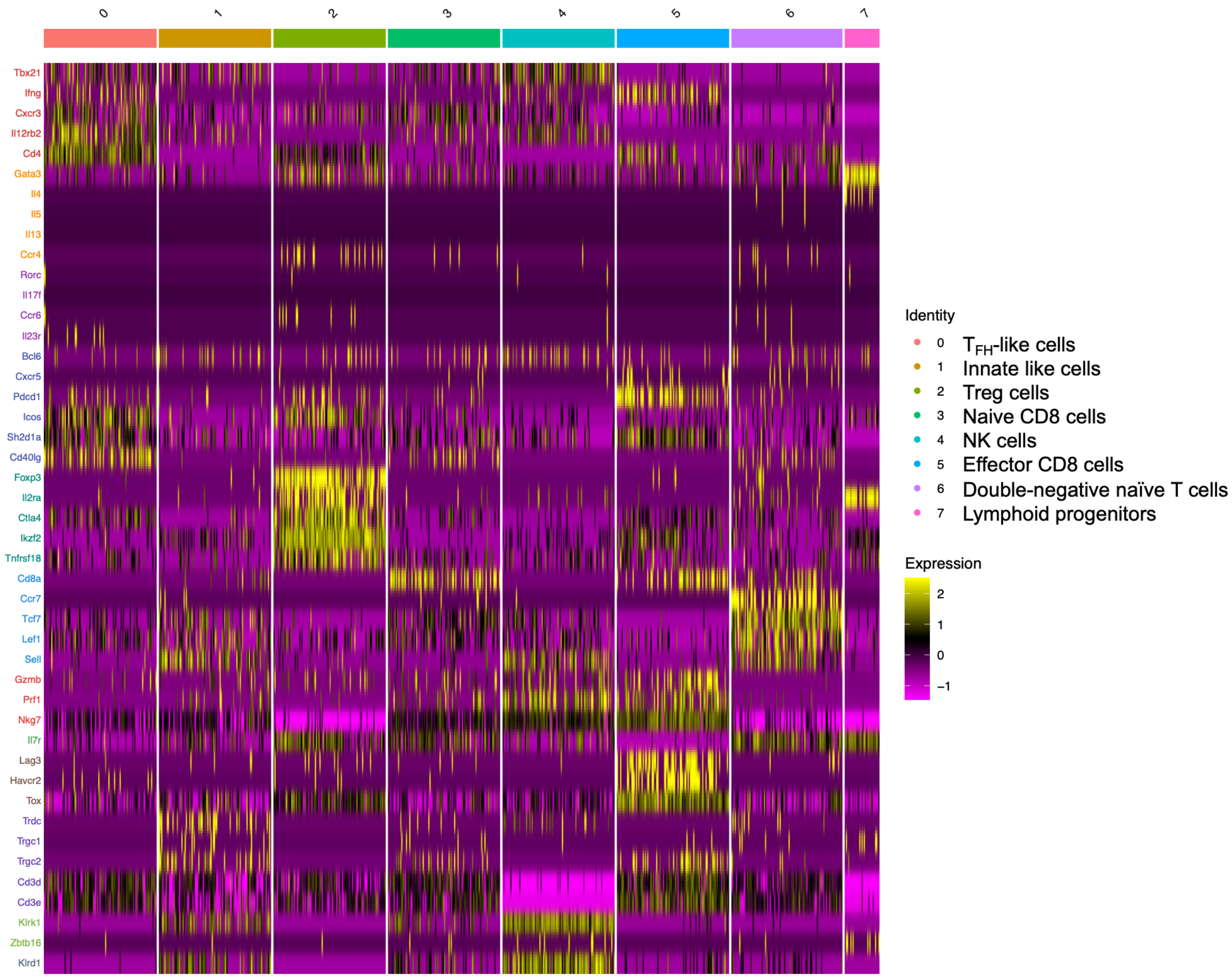


**Supplemental Figure 6. Marker gene expression of indicated T cell subtypes.** Data from further analyses of T-cell subpopulations of supplemental Figure 5.

### Supplemental Table 1

Antibodies for flow cytometry

| Antigen name | Clone | Brand | Catalog number | Fluorescence |
|---|---|---|---|---|
| mouse CD11b | M1/70 | Invitrogen | 17-0112-83 | APC |
| mouse Ly6G | 1A8 | BioLegend | 127607 | PE |
| mouse Ly6C | HK1.4 | BioLegend | 128031 | BV421 |
| mouse c-Kit | 2B8 | BioLegend | 105823 | PerCP-Cy5.5 |
| mouse CD45 | 30-F11 | BD | 564279 | BUV395 |
| mouse CD3e | 145-2C11 | BD | 564379 | BV786 |
| mouse CD8a | 53-6.7 | BioLegend | 104008 | APC-Fire™ 810 |
| mouse CD25 | 3C7 | BD | 570767 | RB780 |
| mouse CD4 | GK1.5 | BD | 571949 | RB670 |
| mouse Foxp3 | MF23 | BD | 560403 | AF488 |
| mouse T-bet | 4B10 | BioLegend | 644814 | APC |
| mouse ICOS | 7E.17G9 | BioLegend | 117429 | BV421 |
| mouse PD-1 | RMP1-30 | BD | 748267 | BV605 |
| mouse CXCR5 | 2G8 (biotinylated) | BD + BioLegend | 551960 + 405241 | SA-BV711 |
| mouse CD40L | SA047C3 | BioLegend | 157014 | APC-Fire™ 750 |

### Supplemental Table 2

Antibodies for immunofluorescence staining

| Target of detection | Primary antibody / dye | Secondary antibody |
|---|---|---|
| mouse Gsdmd | Anti-mouse Gsdmd, Invitrogen PA5-115330, Rabbit (1:200) | Goat anti-Rabbit IgG (H+L) Cross-Adsorbed Secondary Antibody (Alexa Fluor® 647) |
| Endothelial cells | biotin-UEA1 ,Vector Lab, B0-1065-2 (1:300) | Streptavidin, Alexa Fluor® 568 conjugate |
| Stromal cells | Anti-mouse CD271, Invitrogen 14-9400-82 (1:250) | Goat anti-Mouse IgG (H+L) Cross-Adsorbed Secondary Antibody (Alexa Fluor® 488) |
| Human CD40L | Anti-human CD40L, Abcam ab303610 | Goat anti-Rabbit IgG (H+L) Cross-Adsorbed Secondary Antibody (Alexa Fluor® 568) |
| Human CD3 | Anti-human CD3 antibody, Abcam ab17143 | Goat anti-Mouse IgG (H+L) Cross-Adsorbed Secondary Antibody (Alexa Fluor® 488) |
